# A First-In-Class Broad Spectrum Inhibitor of Copper Exporting P_1B_-type ATPases

**DOI:** 10.64898/2026.01.22.700703

**Authors:** Vinit C. Shanbhag, Samuel Anakpeba-Dinguyella, Nikita Gudekar, Kristyn Conrad, Chiemerie Azubuogu, Corinna Probst, Martina Ralle, María G. Mediavilla, Julia A.Cricco, Natalie M. Garza, Vishal M. Gohil, Scott Peck, Siddhartha Kumar, Amarnath Natarajan, Madujika A. Horadigala-Gamage, Gabriele Meloni, Kamal Singh, Michael J. Petris

## Abstract

Copper (Cu) transporting ATPases represent a highly conserved subclass of P-type ATPases with critical roles in Cu export and metalloenzyme synthesis. Despite their important biological roles and association with a wide range of human diseases, no high-affinity small-molecule inhibitors have been described. Here, we identify MKV3 as a first-in-class inhibitor of Cu-transporting P-type ATPases that targets a conserved Cu^+^ entry site to the translocation pathway. *In silico* docking against the *Xenopus* ATP7B structure revealed a highly conserved pocket suitable for pharmacological inhibition. MKV3 bound human ATP7A and ATP7B with nanomolar affinity, competed with N-terminal metal-binding domains for access to the Cu^+^ entry site, and selectively inhibited *Escherichia coli* CopA ATPase activity and Cu^+^ transport. Mechanistically, MKV3 blocked chaperone-mediated Cu^+^ delivery to the intramembranous CPC site of CopA that is essential for its transport function. We further identified a single charged P-domain residue that governed MKV3 affinity and potency across species. Functionally, MKV3 phenocopied the genetic loss of Cu^+^-ATPases in bacteria, fungi, plants, zebrafish, and mammals, impairing copper-dependent enzymes, transporter trafficking, and copper tolerance. These findings establish a conserved, druggable vulnerability in Cu^+^-ATPases and introduce MKV3 as a broadly active chemical tool to modulate copper homeostasis across biological kingdoms.

**Significance Statement:** Copper-transporting P_1B_-type ATPases are essential for copper homeostasis in all domains of life, yet have lacked pharmacological inhibitors. This work identifies MKV3 as the first small-molecule inhibitor of Cu^+^-ATPases in bacteria, fungi, plants and animals, and defines a conserved, druggable Cu^+^ entry pocket that governs metal delivery to the transmembrane pathway. MKV3’s ability to potentiate copper-mediated killing in multidrug-resistant bacterial pathogens highlights its potential as an antimicrobial adjuvant, while its attenuation of mammalian ATP7A/B function offers promise in oncology and copper-related diseases. Collectively, these findings establish a new tool for targeting of Cu^+^-ATPases with wide-ranging applications across biological systems.

## Introduction

Copper (Cu) is an essential micronutrient required by virtually all forms of life, yet it becomes highly toxic when present in excess. Its ability to alternate between reduced (Cu^+^) and oxidized (Cu^2+^) states underlies its role as a catalytic cofactor in numerous enzymes that drive fundamental metabolic processes (1–3). When present at supraphysiological levels, however, copper exerts cytotoxic effects through several mechanisms, including the generation of reactive oxygen species and the displacement of other essential metals from metalloenzymes (4, 5). Excess copper can also trigger cuproptosis, a recently defined form of regulated cell death characterized by copper-dependent aggregation of lipoylated enzymes of the citric acid cycle (6). For thousands of years, humans have exploited the biocidal properties of copper to maintain potable water and prevent wound infections (7). In the modern era, copper is incorporated into wound dressings and frequently touched surfaces in healthcare settings to limit nosocomial infections (8), and it remains widely used as a fungicidal and algicidal agent in agricultural applications (9).

To support essential biochemical functions while avoiding toxicity, organisms have evolved highly coordinated homeostatic systems that tightly regulate copper uptake, intracellular distribution, and export (1). Central to these systems are copper-exporting P-type ATPases (Cu^+^-ATPases)(10), which bioinformatic analyses suggest are present in virtually all organisms (11). Cu^+^-ATPases belong to the P_1B_ subclass of transition-metal-transporting P-type ATPases, which couple ATP hydrolysis to ATP-dependent transmembrane metal ion transport. Within this subclass, Cu^+^-ATPases comprise the P_1B-1_ and P_1B-3_ branches (12). In prokaryotes, Cu^+^-ATPases function primarily in copper detoxification by exporting excess Cu^+^ across the plasma membrane. In contrast, eukaryotic Cu^+^-ATPases have evolved additional specialized roles, including the targeted delivery of copper to nascent cuproenzymes within the secretory pathway. The two human Cu^+^-ATPases, ATP7A and ATP7B, supply copper to cuproenzymes in the *trans*-Golgi network under low-copper conditions, but relocalize to post-Golgi compartments, such as the lysosome or plasma membrane, to facilitate copper efflux when cytosolic copper levels rise (13, 14). Pathogenic variants in ATP7A and ATP7B disrupt these processes and cause lethal disorders of copper metabolism, Menkes disease and Wilson disease, respectively (15). Beyond these genetic disorders, the medical importance of ATP7A and ATP7B is underscored by their documented roles in tumor growth, metastasis, and resistance to platinum-based chemotherapies (16–20). Cu^+^-ATPases are also of broad relevance across diverse biological systems. In plants, these transporters are essential for photosynthesis, ethylene signaling, and copper allocation to developing seeds (21–25). In microbial pathogens, Cu^+^-ATPases function as key virulence factors that mitigate copper stress imposed by the host immune defenses, highlighting copper transport as a conserved determinant of pathogen fitness (26–28). Despite their widespread roles in physiology and disease, Cu^+^-ATPases remain largely unexplored as pharmacological targets, and direct inhibitors of this transporter subclass have not been described.

Like all P-type ATPases, Cu^+^-ATPases operate via the Post-Albers catalytic cycle, in which ATP hydrolysis drives interconversion between discrete conformational states (E1, E1P, E2, and E2P) through auto-phosphorylation and subsequent auto-dephosphorylation of a conserved aspartate residue (29, 30). The catalytic cycle is coordinated by three domains common to all P-type ATPases including a nucleotide-binding (N) domain, a phosphorylation (P) domain and an actuator (A) domain (Fig. 1A). Cu^+^-ATPases contain eight transmembrane helices that form the membrane (M) domain, including six core helices (M1-M6) and two class-specific helices, MA and MB (Fig. 1A). The M4 helix contains a signature CPC motif required for intramembranous Cu^+^-binding during translocation (10). Additional conserved motifs implicated in Cu^+^ coordination and translocation include a YN motif in M5 and an MxxxS motif in M6 (Fig. 1A). P_1B-1_-type Cu^+^-ATPases also harbor variable numbers of N-terminal metal-binding domains (MBDs) which increase with organismal complexity, ranging from one or two in bacteria to six in vertebrates (31). Structural analyses of Cu^+^-ATPases from bacteria, plants, amphibians and humans reveal a highly conserved overall architecture (31–35), including a proposed Cu^+^ entry site formed by an amphipathic “platform” generated by a bend in the MB-MB’ helix as it emerges from the lipid bilayer into the cytoplasm (Fig. 1A). This platform lies adjacent to a conserved M-E-D triad, comprising methionine, glutamate, and aspartate residues at the cytoplasmic interfaces of M1, M2, and M3, respectively, which are predicted to facilitate Cu^+^ entry (Fig. 1A). Biochemical, biophysical, and structural studies support a three-stage model for Cu^+^ translocation involving: (i) cytoplasmic Cu^+^ entry mediated by docking of a copper metallochaperone or an N-terminal MBD to the platform; (ii) Cu^+^ transfer to the intramembrane CPC site facilitated by transient interactions with the methionine residue in the M-E-D triad; and (iii) Cu^+^ release through an exofacial exit pathway (36).

**Figure 1:**
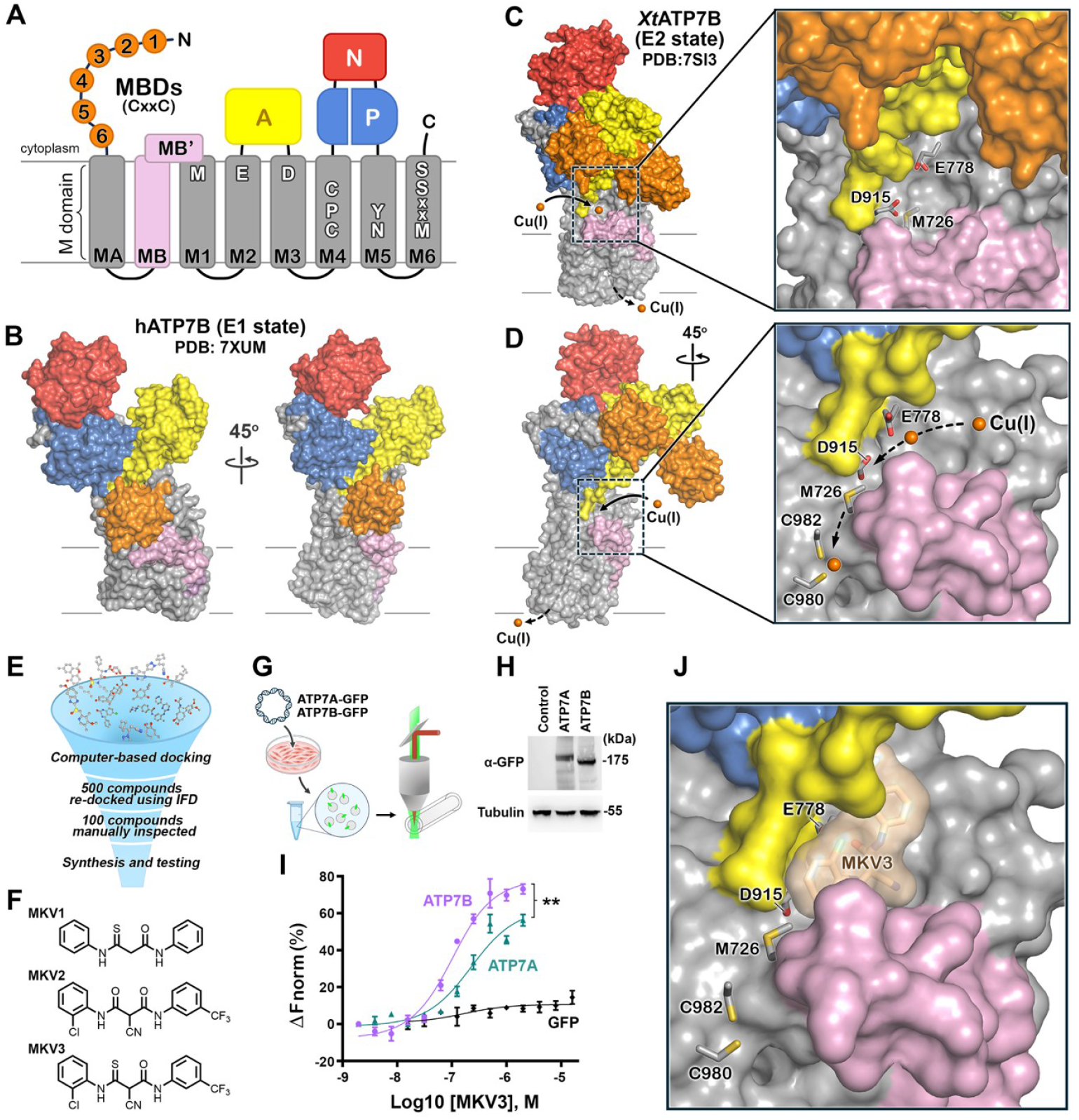
Rational design of small molecule inhibitors targeting P_1B_-type copper ATPases. (A) Schematic representation of Cu^+^-ATPases showing eight transmembrane domains (MA and M1–M6, gray) and the MB/MB′ platform (pink). Signature motifs include the conserved M–E–D triad (M1–M3), Cu^+^-binding CPC motif (M4), YN motif (M5), and MxxxS motif (M6). The N-terminal metal-binding domains (MBDs, orange) contain Cu^+^-binding CxxC motifs, and the cytoplasmic catalytic domains include the actuator (A, yellow; TGE motif), nucleotide-binding (N, red), and phosphorylation (P, blue; DKTGT motif) domains. (B) Surface rendering of human ATP7B in the E1 state (hATP7B; PDB: 7XUM) shown in two orientations. (C) Surface structure of *Xenopus tropicalis* ATP7B (*Xt*ATP7B; PDB ID: 7SI3) shown in the E2 state, highlighting the Cu(I) entry site and the conserved M–E–D triad (M726, E778, D915). (D) A 45° rotated view of *Xt*ATP7B showing the spatial relationship between the M–E–D triad and the downstream CPC motif (C980, C982) that lines the Cu(I) translocation pathway (inset). (E) A schematic workflow summarizes the small-molecule screen used to identify inhibitors. (F) Chemical structures of representative hit compounds (MKV1, MKV2, and MKV3). (G) The experimental workflow for Microscale Thermophoresis (MST) analysis is shown, where detergent-free lysates from HEK293 cells expressing GFP-tagged ATP7A/7B (and GFP control) were analyzed for MKV3 binding. (H) Immunoblot confirming expression of ATP7A-GFP and ATP7B-GFP in lysates used for MST analysis. (I) MST binding curves showing concentration-dependent interaction of MKV3 with ATP7A and ATP7B, with no binding detected in GFP-only controls. (J) Surface rendering of *Xt*ATP7B with MKV3 docked in the Cu-entry pocket (best-scoring pose, −6.99 kcal/mol) is shown. Data are represented as mean ± SEM; **p < 0.01.

Here, we describe the discovery of MKV3, a first-in-class small-molecule inhibitor of Cu^+^-ATPases, identified through a structure-guided *in-silico* screen targeting a conserved Cu^+^ entry pocket and active across bacteria, fungi, protists, plants, and animals. Biochemical analyses suggest that MKV3 binds within the Cu^+^ entry pocket, thereby blocking Cu^+^ delivery to the intramembranous transport site. Mutagenesis studies identify a single charged P-domain residue at the rear of the Cu^+^ entry pocket which governs MKV3 affinity and potency across species.

Together, these findings reveal an evolutionarily conserved vulnerability in P_1B_-type ATPases and establish a chemical tool for targeting copper transport in diverse biological systems.

## Results

### Computational screening identifies candidate inhibitors of the Cu entry pocket

To identify potential sites for pharmacological inhibition, we first examined the published structure of the *Xenopus tropicalis* ATP7B (*Xt*ATP7B) ortholog in its E2P conformation (PDB 7SI3) (34). This structure reveals a cavity adjacent to the platform domain through which Cu^+^ is predicted to enter the transmembrane domain and subsequently directed toward the intramembranous CPC motif by methionine, glutamate and aspartate residues of the M-E-D triad (Fig. 1A, 1C, and 1D). Notably, comparison with the human ATP7B structure in the E1 conformation (Fig. 1B), reveals this Cu^+^ entry pocket is occupied by the fifth N-terminal metal-binding domain (MBD5), supporting a model in which this pocket serves as a docking site for Cu^+^ delivery to the transport pathway (33). Structural modelling of Cu^+^-ATPases in the E2 conformation across bacteria, fungi, plants, and metazoans revealed a striking preservation of the M-E-D triad and surrounding pocket architecture (Fig. S1 and S2) and suggested that a small molecule designed to occupy this site could, in principle, exhibit broad inhibitory activity across diverse species.

Next, we employed an *in-silico* docking strategy to identify compounds capable of binding within the Cu^+^ entry pocket of XtATP7B. A curated chemical library was screened using the Glide docking algorithm within the Schrödinger software suite, and hit compounds were ranked based on predicted binding energy (Fig. 1E) (37, 38). Three top-scoring candidates, designated MKV1, MKV2, and MKV3, with a common molecular scaffold were selected for synthesis and further experimental testing (Fig. 1F). To experimentally assess compound binding to Cu^+^-ATPases, we employed microscale thermophoresis (MST), a sensitive biophysical technique capable of detecting ligand interactions with GFP-tagged membrane proteins in detergent-free whole cell lysates (39). HEK293 cells were transiently transfected with plasmids encoding either GFP alone or GFP-tagged human ATP7A or ATP7B, and lysates including membrane vesicles were prepared by sonication under detergent-free conditions (Fig. 1G). The expression of ATP7A and ATP7B in HEK293 lysates was confirmed by immunoblotting (Fig. 1H). MST analysis revealed no detectable binding of any compound to GFP alone. There was a clear, dose-dependent binding of MKV3 to GFP-ATP7A with a dissociation constant (K_d_) of 219 ± 14 nM (Fig. 1I). In contrast, MKV1 exhibited a significantly lower affinity while there was no binding detected for MKV2 (Table S1). In parallel studies, MKV3 exhibited dose-dependent binding to GFP-tagged human ATP7B with an even higher affinity (K_d_, 69.5 ± 6.5 nM) (Fig. 1I). Based on these results, MKV3 was selected as our lead candidate for further studies. A docking model of MKV3 within the Cu entry pocket of *Xt*ATP7B is shown (Fig. 1J).

### Metal-binding domains, MBD5 and MBD6, compete with MKV3 for binding to human ATP7A

Since the predicted MKV3 binding pocket in the E2P conformation of *Xt*ATP7B overlaps with the site occupied by either MBD5 or MBD6 in the E1 structure of human ATP7B (Fig. 2A and 2B), we hypothesized that these metal-binding domains might compete with MKV3 for access to the Cu^+^ entry site. To test this, we performed MST analysis of MKV3 binding to truncated GFP-ATP7A variants that differed in their MBD5/6 composition as follows: (i) GFP-ATP7A lacking the first 422 amino acids encompassing MBDs1-4 leaving MBD5 and MBD6 intact (WT-MBD5/6); (ii) a WT-MBD5/6 variant in which MBD5 and MBD6 were deleted (ΔMBD5/6); and iii) a WT-MBD5/6 variant in which the Cu-coordinating cysteines of both MBD5/6 were replaced with serines (mMBD5/6) (Fig. 2C). Each construct was transiently expressed in HEK293 cells, and immunoblot analysis confirmed comparable expression levels across samples used for MST measurements (Fig. 2D). MST analysis suggested MKV3 bound WT-MBD5/6 with a K_d_ of 162 ± 7 nM, whereas binding affinity increased nearly ten-fold in the ΔMBD5/6 variant (K_d_ = 18 ± 3 nM). Substitution of the CxxC motifs with serines partially restored affinity (K_d_ = 55 ± 5 nM), indicating that disruption of Cu-binding capacity reduces, but does not eliminate, competition with MKV3 (Fig. 2E and 2F). These results are consistent with a model in which MKV3 binds within the Cu^+^ entry pocket occupied by MBD5/6 to deliver Cu^+^ to the transmembrane transport pathway.

**Figure 2.**
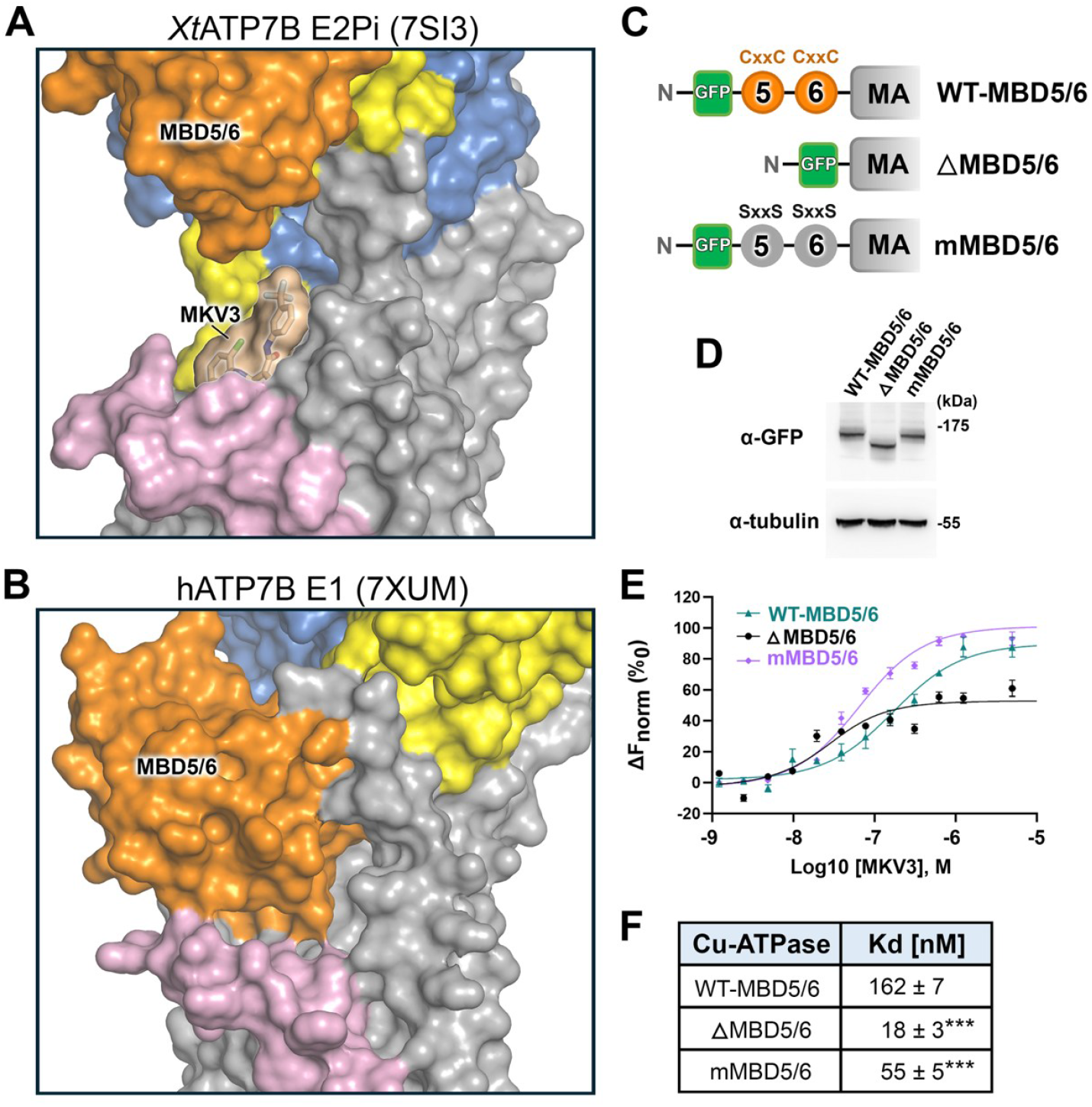
Loss of the N-terminal metal binding domains enhances MKV3 binding to ATP7A. (A) Surface representation of the E2-Pi conformation of *Xt*ATP7B (PDB ID: 7SI3), showing displacement of MBD5/6, which increases accessibility of the copper entry site. MKV3 shown in surface view is highlighted to show its binding within the pocket. (B) Surface representation of the E1 state of human ATP7B (PDB ID: 7XUM), showing MBD5/6 (orange) positioned near the copper entry site. (C) Schematic representation of GFP-tagged ATP7A constructs: WT-MBD5/6 (truncated version without the MBD1-4), ΔMBD5/6 (lacking MBD5 and MBD6), and mMBD5/6 (with CxxC motifs replaced by SxxS in MBD5 and MBD6). (D) Immunoblot analysis of WT-MBD5/6, ΔMBD5/6, and mMBD5/6 constructs expressed in HEK293 cells, showing similar protein levels normalized to tubulin. (E) MST analysis of MKV3 binding to WT-MBD5/6, ΔMBD5/6, and mMBD5/6 ATP7A variants. (F) MKV3 binding affinities (mean K_d_ ± SEM), showing significantly stronger binding to ΔMBD5/6 and mMBD5/6 compared to WT-MBD5/6 (Data are represented as mean ± SEM; ***p < 0.001).

### MKV3 selectively inhibits the *Escherichia coli* Cu^+^-ATPase

To determine whether MKV3 functions as an inhibitor of Cu^+^-ATPases, we examined its effect on the *Escherichia coli* Cu^+^-ATPase CopA (*Ec*CopA). We have previously characterized the ATPase and Cu^+^ transport activities of purified recombinant *Ec*CopA reconstituted in proteoliposomes (40). Using this system, we observed a dose-dependent decrease in Cu^+^-stimulated ATPase activity in the presence of MKV3 (Fig. 3A and 3C) with an IC_50_ of 130 ± 19 nM. In contrast, the MKV2 compound, which failed to bind ATP7A, showed no detectable inhibition. To assess specificity, we tested the effect of MKV3 on the ATPase activity of *Ec*ZntA, a P_1B-2_-type ATPase that transports Zn^2+^, Cd^2+^, and Pb^2+^ (Fig. 3B). MKV3 had no measurable effect on Zn^2+^-stimulated *Ec*ZntA ATPase activity, and MKV2 also showed no inhibition. These results suggest that MKV3 selectively targets the Cu^+^-specific P_1B-1_-type ATPase *Ec*CopA without affecting related transition-metal pumps.

**Figure 3.**
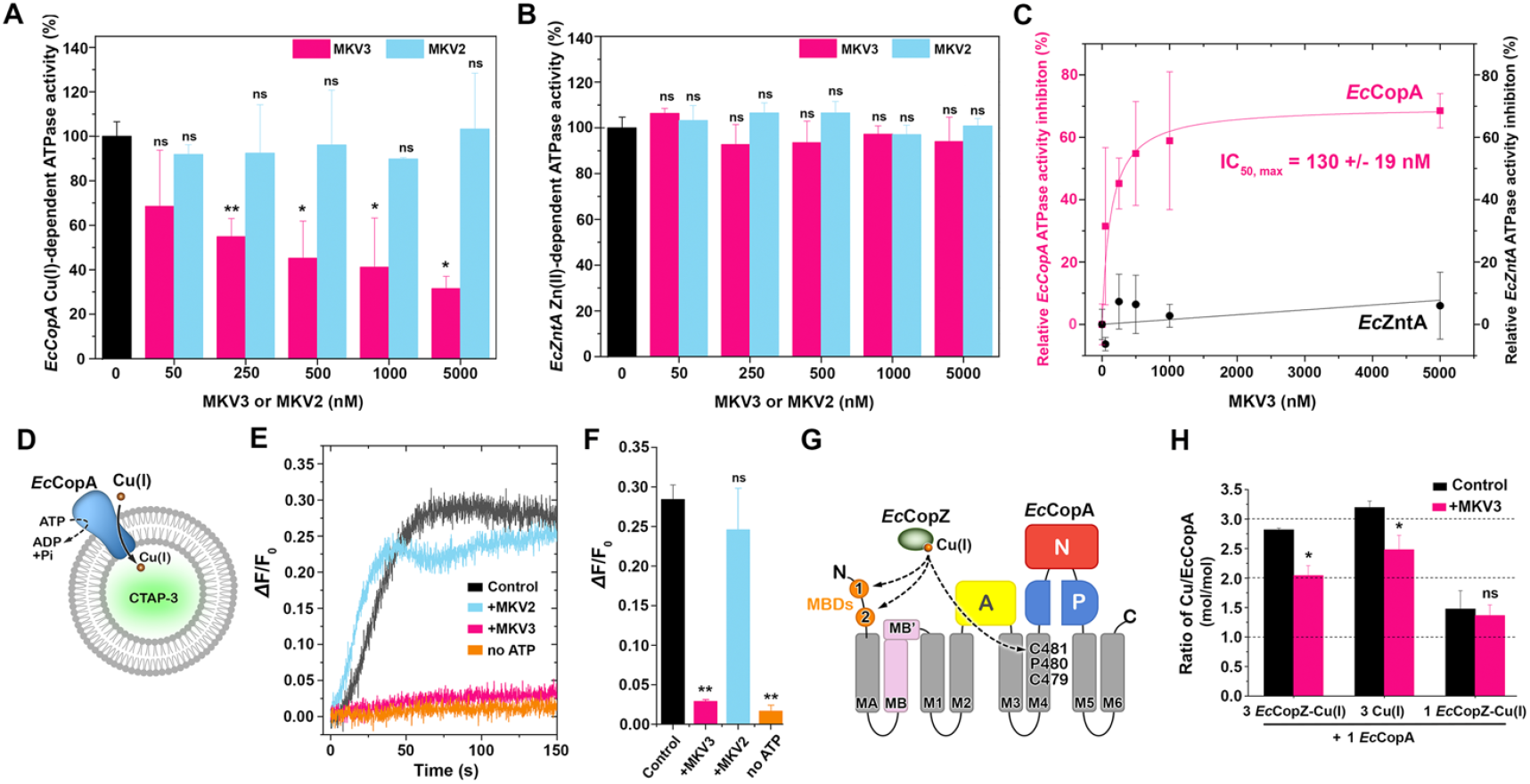
MKV3 inhibits *E. coli* CopA function by blocking chaperone-mediated Cu(I) delivery. (A) Cu(I)-stimulated ATPase activity of *Ec*CopA in the presence of varying concentrations of MKV3 (IC_50_ = 130 ± 19 nM). (B) Zn(II)-stimulated ATPase activity of *Ec*ZntA in the presence of MKV3, used as a control for specificity. (C) Dose-response curves showing the effects of MKV3 on ATPase activities of *Ec*CopA and *Ec*ZntA. Data are normalized to percentage ATPase activity inhibition (D) Proteoliposome assay to measure Cu(I) transport by *Ec*CopA using the Cu-responsive fluorescent probe CTAP-3. (E) Real-time CTAP-3 fluorescence emission kinetic traces of Cu(I) transport by *Ec*CopA in the presence or absence of MKV3, including MKV2 and no-ATP controls. (F) Quantification of CTAP-3 fluorescence emission changes from Cu(I) transport assays under the indicated conditions. (G) Schematic representation of *Ec*CopZ-mediated Cu(I) delivery to *Ec*CopA, showing copper transfer to the N-terminal metal-binding domains (MBDs) and the transmembrane CPC motif. (H) Quantification of copper transfer from *Ec*CopZ to *Ec*CopA in the presence or absence of MKV3, as determined by ICP-MS. Data are represented as mean ± SEM; *p < 0.05, **p < 0.01, ns = not significant.

To determine whether MKV3 also blocks Cu^+^ transport, we measured its effect on *Ec*CopA-dependent Cu^+^ translocation into proteoliposomes containing in their lumen CTAP-3, a water-soluble and membrane-impermeant fluorescent Cu^+^ probe (Fig. 3D). Addition of Cu^+^ to CTAP-3-encapsulated *Ec*CopA proteoliposomes produced a time-dependent increase in fluorescence, indicating Cu^+^ translocation across the lipid bilayer (Fig. 3E and 3F). No fluorescence increase was detected in reactions lacking ATP, confirming this energy source requirement for active primary transport. When MKV3 was included, no fluorescence signal increase over time was observed, demonstrating complete inhibition of Cu^+^ translocation. MKV2 had no effect under the same conditions. These findings indicate that MKV3 specifically inhibits *Ec*CopA-mediated ATP hydrolysis and Cu^+^ transport, consistent with a direct blockade of the Cu^+^ entry pathway.

### MKV3 inhibits metallochaperone-mediated Cu^+^ delivery to *Ec*CopA

We next tested whether MKV3 interferes with Cu^+^ transfer from the metallochaperone *Ec*CopZ to its cognate Cu^+^-ATPase *Ec*CopA. The delivery of Cu^+^ from CopZ homologs to CopA-type transporters has been well characterized (41–43). Full-length *Ec*CopA contains three high-affinity Cu^+^-binding sites: two located within its N-terminal MBDs and one within the transmembrane CPC motif (Fig. 3G). If MKV3 binds within the Cu^+^ entry pocket, it should block Cu^+^ transfer from *Ec*CopZ to the transmembrane site but not to the N-terminal MBDs. To test this hypothesis, purified *Ec*CopA was incubated with Cu^+^-loaded *Ec*CopZ (*Ec*CopZ-Cu^+^) at 1:3 and 1:1 molar ratios under anaerobic conditions in the presence or absence of stoichiometric MKV3 (Fig. 3H). A control reaction containing a three-fold molar excess of free Cu^+^ was included to confirm maximal metalation. After incubation, *Ec*CopA was separated by size-exclusion chromatography, and Cu^+^ content was quantified as a function of *Ec*CopA concentration. In the absence of MKV3, both *Ec*CopZ-Cu^+^ and free Cu^+^ produced stoichiometric metalation of three Cu^+^-binding sites within *Ec*CopA. When MKV3 was added, the number of Cu^+^ ions bound per *Ec*CopA molecule decreased to two, consistent with inhibition of Cu^+^ transfer to the transmembrane CPC site (Fig. 3H). MKV3 did not reduce Cu^+^ transfer when sub-stoichiometric amounts of *Ec*CopZ-Cu^+^ were used, suggesting that MKV3 does not interfere with Cu^+^ loading of the N-terminal MBDs. These findings suggest MKV3 prevents Cu^+^ delivery from *Ec*CopZ to the intramembranous CPC site without disrupting metalation of the N-terminal domains.

### MKV3 inhibits ATP7A-dependent copper transport in mammals and zebrafish

We next examined whether MKV3 inhibits ATP7A, the principal Cu^+^-ATPase in mammals. Under basal conditions, ATP7A localizes to the *trans*-Golgi network (TGN), where it delivers copper to nascent cuproenzymes within the secretory pathway. When cytosolic copper increases, ATP7A traffics from the TGN to post-Golgi vesicles and the plasma membrane to mediate copper efflux and maintain homeostasis (Fig. 4A). Notably, this trafficking requires catalytic activity of the transporter (44). Therefore, a *bona fide* ATP7A inhibitor should (i) reduce activity of copper-dependent secretory enzymes, (ii) prevent copper-stimulated trafficking, (iii) increase cellular copper levels due to impaired efflux, and (iv) enhance copper sensitivity.

**Figure 4.**
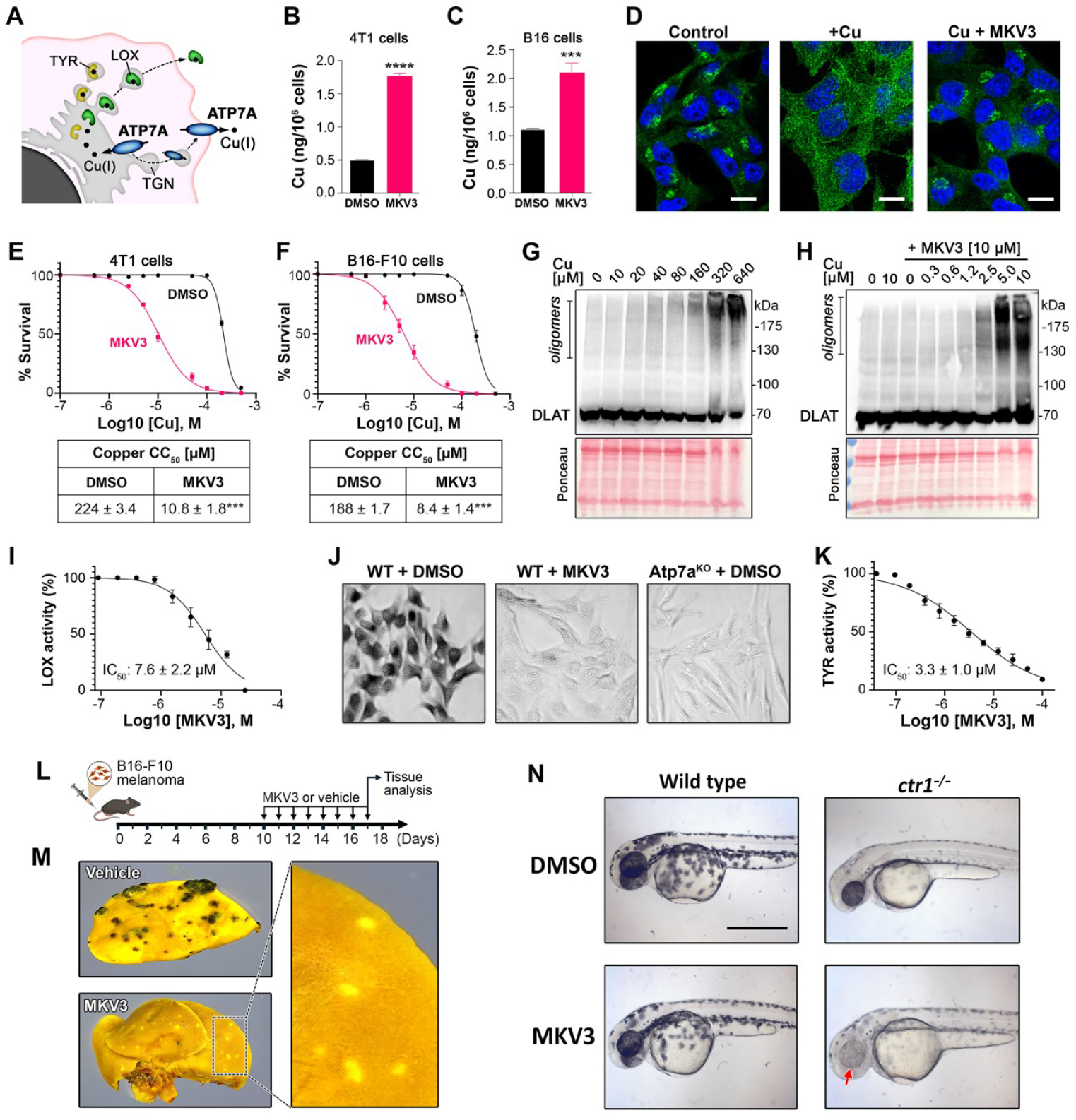
MKV3 Inhibits ATP7A-dependent copper transport in vertebrates. (A) Schematic illustration of ATP7A function in mammalian cells. ATP7A resides within the TGN, delivering Cu to cuproenzymes in the secretory pathway, including tyrosinase (TYR) and lysyl oxidase (LOX). Elevated Cu concentrations stimulate ATP7A trafficking to post-Golgi vesicles and the plasma membrane for Cu efflux. (B, C) Total cellular copper concentrations in 4T1 and B16 cells treated for 24 hours with or without 10 µM MKV3, measured using ICP-MS. (D) ATP7A immunofluorescence in 4T1 cells treated with Cu and MKV3. Cells were pretreated with 10 µM BCS for 16 hours, then exposed for 4 hours to 2 µM Cu alone, 2 µM Cu with 10 µM MKV3, or 1% DMSO vehicle control. ATP7A was detected by immunofluorescence (green), and nuclei were stained with DAPI (blue). Scale bars, 10 µm. (E, F) Survival of 4T1 and B16 cells after a 48-hour treatment with the indicated Cu concentrations with or without 10 µM MKV3. Copper CC_50_ values: 4T1 cells (DMSO = 224 ± 3.4 µM Cu; MKV3 = 10.8 ± 1.8 µM Cu); B16 cells (DMSO = 188 ± 1.7 µM Cu; MKV3 = 8.4 ± 1.4 µM Cu). (G) MKV3 enhances Cu-mediated cuproptosis. Immunoblot analysis of DLAT oligomerization in 4T1 cells treated with Cu alone or (H) Cu plus MKV3 at indicated concentrations. (I) MKV3 inhibits ATP7A-dependent LOX activity. LOX activity in the media of 4T1 cells treated for 5 days with or without MKV3 (IC_50_ = 7.6 ± 2.2 µM). (J) MKV3 inhibits ATP7A-dependent tyrosinase activity. *In situ* tyrosinase activity in wild-type B16 cells (WT) treated with DMSO or 5 µM MKV3 for 24 hours, alongside knockout Atp7a^KO^ B16 cells (DMSO) serving as a control for ATP7A-dependent pigmentation. (K) Tyrosinase activity in wild-type B16 cells treated for 24 hours with the indicated concentrations of MKV3 (IC_50_ = 3.3 ± 1.0 µM). (L) Experimental timeline showing B16-F10 melanoma implantation, MKV3 or vehicle dosing, and tissue analysis. (M) B16-F10 tumor-bearing mice received daily subcutaneous injections of MKV3 (50 mg/kg) or vehicle (5% DMSO /5% Tween-80) for 7 days. Lungs were fixed, stained with Bouin’s solution, and imaged. (N) MKV3 reduces melanogenesis in zebrafish. WT and *ctr1*^-/-^ zebrafish embryos at 3 hours post-fertilization were treated for 48 hours with or without 1 µM MKV3. Note the reduced pigmentation in the *ctr1*^-/-^ embryo (red arrow).

To test these predictions, we first measured cellular copper levels in 4T1 breast cancer and B16-F10 melanoma cells treated for 24 h with 10 μM MKV3, a concentration well below cytotoxic levels in these cells (Fig. S3). Consistent with inhibition of copper export, MKV3 significantly increased cellular copper accumulation in both cell lines (Fig. 4B and 4C). MKV3 also inhibited copper-stimulated trafficking of ATP7A from its perinuclear TGN localization to cytoplasmic vesicles in 4T1 cells (Fig. 4D). Sensitivity to cytotoxic copper was also enhanced by MKV3, reducing the copper CC_50_ by approximately 20-fold in both 4T1 and B16-F10 cells (Fig. 4E and 4F). MKV3 enhanced copper-dependent cell death by cuproptosis, lowering the amount of exogenous copper required to induce oligomerization of mitochondrial DLAT (Fig. 4G and 4H). In addition, MKV3 potentiated cytotoxicity of Ag^+^, an isoelectronic Cu^+^ mimetic and known substrate of P_1B_-type Cu^+^-ATPases (45), while having no effect on Co^2+^, Fe^2+^, Mn^2+^ or Zn^2+^, sensitivity (Fig. S4). Taken together, these results suggest that MKV3 selectively potentiates copper and silver sensitivity, triggering cuproptosis through inhibition of ATP7A trafficking and Cu^+^ efflux.

We next examined whether MKV3 impairs ATP7A-mediated copper delivery to two cuproenzymes, lysyl oxidase and tyrosinase, which receive copper from ATP7A within the secretory pathway (Fig. 4A) (19, 46). Treatment of 4T1 cells with MKV3 significantly reduced the activity of lysyl oxidase in the conditioned medium compared to vehicle control (Fig. 4I). MKV3 significantly reduced the activity of tyrosinase in wild-type B16-F10 cells, phenocopying the loss of tyrosinase activity observed following CRISPR-mediated disruption of the *Atp7a* gene in these cells (Atp7a^KO^) (Fig. 4J and 4K). Notably, Atp7a^KO^ cells transiently transfected with expression plasmids encoding either human ATP7A or ATP7B restored tyrosinase activity, which was suppressed by MKV3 added to the medium, indicating MKV3 can target both mammalian copper ATPases (Fig. S5). In a mouse model of metastatic B16-F10 melanoma, subcutaneous administration of MKV3 suppressed melanogenesis in metastatic lung tumors *in vivo* (Figs. 4L and 4M, and Fig. S6). Collectively, these results demonstrate that MKV3 inhibits ATP7A-dependent copper efflux and copper delivery to cuproenzymes of the secretory pathway.

To test whether MKV3 also inhibits ATP7A homologs in lower vertebrates, we evaluated its effects on melanogenesis in zebrafish embryos. In wild-type embryos, MKV3 at concentrations above 1 μM caused severe developmental defects and lethality whereas non-toxic doses (≤1 μM), failed to impact melanogenesis (Fig. 4N and Fig. S7). To increase sensitivity to copper loss, we used *ctr1*^−^*/*^−^ embryos lacking the high-affinity copper importer CTR1, which display reduced pigmentation due to impaired copper uptake. Treatment of *ctr1*^−^*/*^−^ embryos with sublethal concentrations of MKV3 further reduced melanogenesis compared to vehicle-treated mutants (Fig. 4N and Fig. S7). Taken together, these data suggest that MKV3 inhibits ATP7A-mediated copper transport in mammals and lower vertebrates.

### Identification of a charged P-domain residue that governs MKV3 binding and potency across species

Our MST analysis suggested that MKV3 binds with higher affinity to human ATP7B than to human ATP7A (Fig. 1I). Structural modeling of MKV3 within the Cu^+^ entry pocket of amphibian *Xt*ATP7B reveals a single non-conserved residue predicted to contact the ligand: E1010 in *Xt*ATP7B, E1030 in human ATP7A, and K1013 in human ATP7B (Fig. 5A, 5B and Table S2). This residue, positioned in the P-domain, is predicted to interact electrostatically with a fluorine atom in MKV3. To test whether this residue is a determinant of MKV3 affinity, we substituted E1030 of ATP7A with a lysine. GFP-tagged wild-type and E1030K variants of ATP7A were expressed at comparable levels in HEK293 cells and MKV3 affinity was measured by MST. The ATP7A-E1030K variant exhibited significantly higher affinity for MKV3 relative to ATP7A-WT (Figs. 5C and 5D). To determine whether this substitution also increases MKV3 potency, both WT and E1030K variants of ATP7A were expressed in Cu^s^ cells, a copper-sensitive cell line lacking ATP7A and metallothioneins I and II that is inviable in basal medium but can be propagated in the presence of a copper chelator (47). As expected, Cu^s^ cells expressing similar levels of either the WT or E1030K mutant of ATP7A grew robustly in basal medium, indicating that the mutation did not impair transporter function (Fig. 5E and 5F). Addition of MKV3 to the medium suppressed growth in a concentration-dependent manner for both constructs, but the E1030K variant displayed a markedly lower IC_50_ (8.6 ± 1.9 µM) than WT ATP7A (19.7 ± 1.6 µM; p < 0.001) (Fig. 5G). These data suggest E1030 as a key determinant of MKV3 binding and potency.

**Figure 5.**
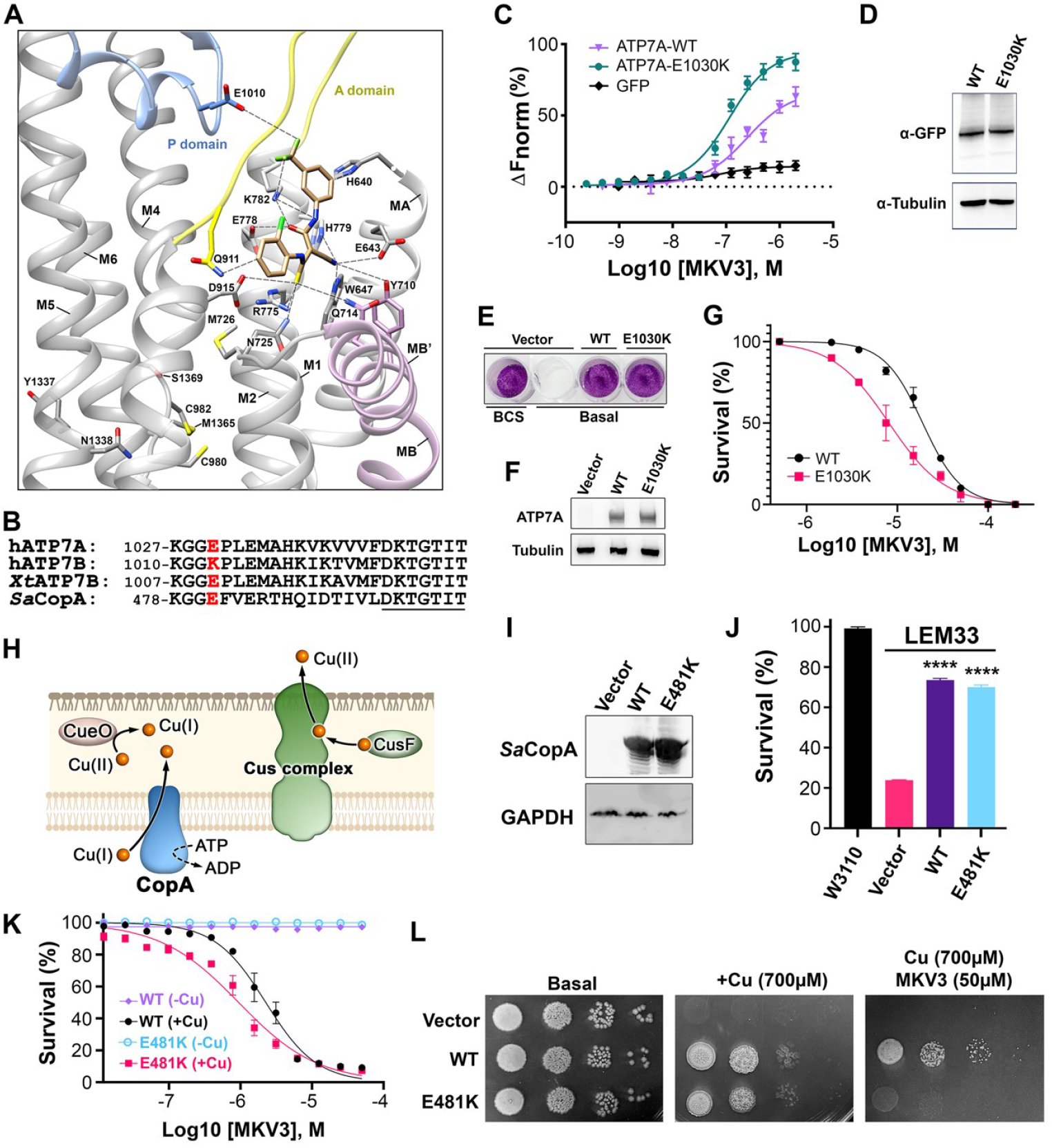
MKV3 potency is determined by a glutamate residue in the binding pocket of ATP7A and *Staphylococcus aureus* CopA. (A) Structural model of MKV3 docked within the Cu^+^ entry pocket of *Xt*ATP7B showing residues predicted to be within 5Å of the ligand (dashed lines). (B) Conservation of amino acids in the predicted MKV3 binding pocket of hATP7A, hATP7B, *Xt*ATP7B, and *Sa*CopA. (C) Microscale thermophoresis analysis of MKV3 binding to GFP-tagged ATP7A-WT and ATP7A-E1030K. Apparent MKV3 binding affinities (mean ± SEM): WT = 178.5 ± 2.5 nM; E1030K = 89.6 ± 4.5 nM; p < 0.01. (D) Immunoblot analysis of HEK293 cells transiently transfected with GFP-tagged WT or E1030K ATP7A. (E) Synthetic lethal phenotype of Cu^S^ cells (ATP7A^−^/^−^; MT^−^) rescued by stable expression of WT or E1030K ATP7A. (F) Immunoblot analysis of ATP7A variants expressed in rescued Cu^S^ cells (WT and E1030K). (G) Survival of Cu^S^ cells expressing WT or E1030K ATP7A after 96-hour exposure to MKV3 at the indicated concentrations. MKV3 CC_50_ values (mean ± SEM): WT = 19.73 ± 1.55 µM; E1030K = 8.61 ± 1.94 µM; p < 0.05. (H) Schematic illustration of *E. coli* copper tolerance pathways deleted in the copper-sensitive LEM33 strain (*ΔcopA, ΔcueO, ΔcusCFBA*). (I) Immunoblot analysis of LEM33 transformed with empty vector or expressing WT or E481K *Sa*CopA. (J) Survival of LEM33 transformed with empty vector, WT *Sa*CopA, or E481K *Sa*CopA relative to the parental *E. coli* W3110 strain in LB medium containing 2 mM Cu. (K) MKV3-mediated copper sensitivity of LEM33 cells expressing WT or E481K *Sa*CopA grown for 16h in LB medium containing the indicated concentrations of MKV3 with or without 2 mM Cu. MKV3 IC_50_ values (mean ± SEM): WT = 2.25 ± 0.47 µM; E481K = 0.94 ± 0.18 µM; p < 0.05. (L) Serial dilutions of LEM33 cells expressing empty vector, WT *Sa*CopA, or E481K *Sa*CopA plated on LB agar containing 700 µM Cu and/or 50 µM MKV3.

To evaluate whether this mechanism is broadly conserved, we examined the CopA Cu^+^-ATPase of *Staphylococcus aureus* (*Sa*CopA), which also contains a glutamate at the analogous P-domain position (E481) (Fig. 5B). Wild-type and E481K variants of *Sa*CopA were expressed in the copper-sensitive *E. coli* strain LEM33 (*ΔcopA, ΔcueO, ΔcusCFBA*), which lacks the endogenous *Ec*CopA transporter, the periplasmic Cu^+^ oxidase CueO, and the Cus system for periplasmic copper export (Fig. 5H) (48). At equivalent expression levels, both wild-type and E481K variants of *Sa*CopA rescued growth of LEM33 cells in LB media containing inhibitory copper concentrations, confirming that the E481K substitution did not compromise Cu^+^ transport (Figs. 5I and 5J). MKV3 suppressed the ability of both variants to complement LEM33 growth in copper-supplemented LB media, however, the E481K mutant was significantly more sensitive compared to wild-type *Sa*CopA (MKV3 IC_50_ = 2.6 ± 0.5 µM vs 0.9 ± 0.2 µM; p < 0.001) (Fig. 5K). This differential potency of MKV3 was readily apparent on agar plates (Fig. 5L). Taken together, these findings identify a single charged residue as a critical determinant of MKV3 affinity and potency across bacterial and mammalian Cu^+^-ATPases.

### MKV3 targets Cu^+^-ATPases across a broad evolutionary distance

To determine the breadth of MKV3 activity across diverse organisms, we tested whether MKV3 can phenocopy loss-of-function mutations in Cu^+^-ATPases from bacteria, fungi, and plants. We first examined a methicillin-resistant *S. aureus* (MRSA) strain (USA300), which is a major cause of community-acquired infections in North America and Europe (49–51). Like all *S. aureus* strains, USA300 expresses the P_1B-1_-type ATPase *Sa*CopA, and its cognate metallochaperone *Sa*CopZ, from the *copAZ* operon (Fig. 6A) (52). In addition, USA300 expresses *Sa*CopB, a Cu^+^-exporting P_1B-3_-type ATPase encoded by the *copBL* operon together with the membrane-associated copper-binding protein CopL (52). Consistent with prior studies (52), growth of the *ΔcopBL;ΔcopAZ* variant of USA300 was abolished on NB1 agar plates containing 1.5 mM CuCl_2_, whereas wild-type, *ΔcopBL* and *ΔcopAZ* variants exhibited robust growth under these conditions (Fig. S8). We next evaluated the ability of MKV3 to create a zone of growth inhibition on copper-supplemented agar. MKV3 generated two distinct zones of growth inhibition for the wild-type USA300 strain, consistent with inhibition of both *Sa*CopA and *Sa*CopB transporters (Fig. 6B and 6C). In contrast, the *ΔcopAZ* and *ΔcopBL* mutant strains each displayed a single, enlarged zone of growth inhibition, suggesting enhanced sensitivity to MKV3 when one of the copper pumps was absent. MKV3 did not affect the growth of any strain on plates lacking supplemental copper, indicating that its antibacterial activity is copper dependent (Fig. S9). Since solid copper surfaces possess innate antimicrobial properties (8), we tested whether MKV3 enhances copper-mediated contact killing of MRSA. Compared to vehicle control, MKV3 significantly enhanced copper-mediated contact killing of the wild-type USA300 strain in a time-dependent manner compared to a stainless-steel control surface (Figs. 6D and 6E; Fig. S10). Taken together, these data suggest MKV3 targets both *Sa*CopA and *Sa*CopB to sensitize MRSA to the antimicrobial properties of copper.

**Figure 6.**
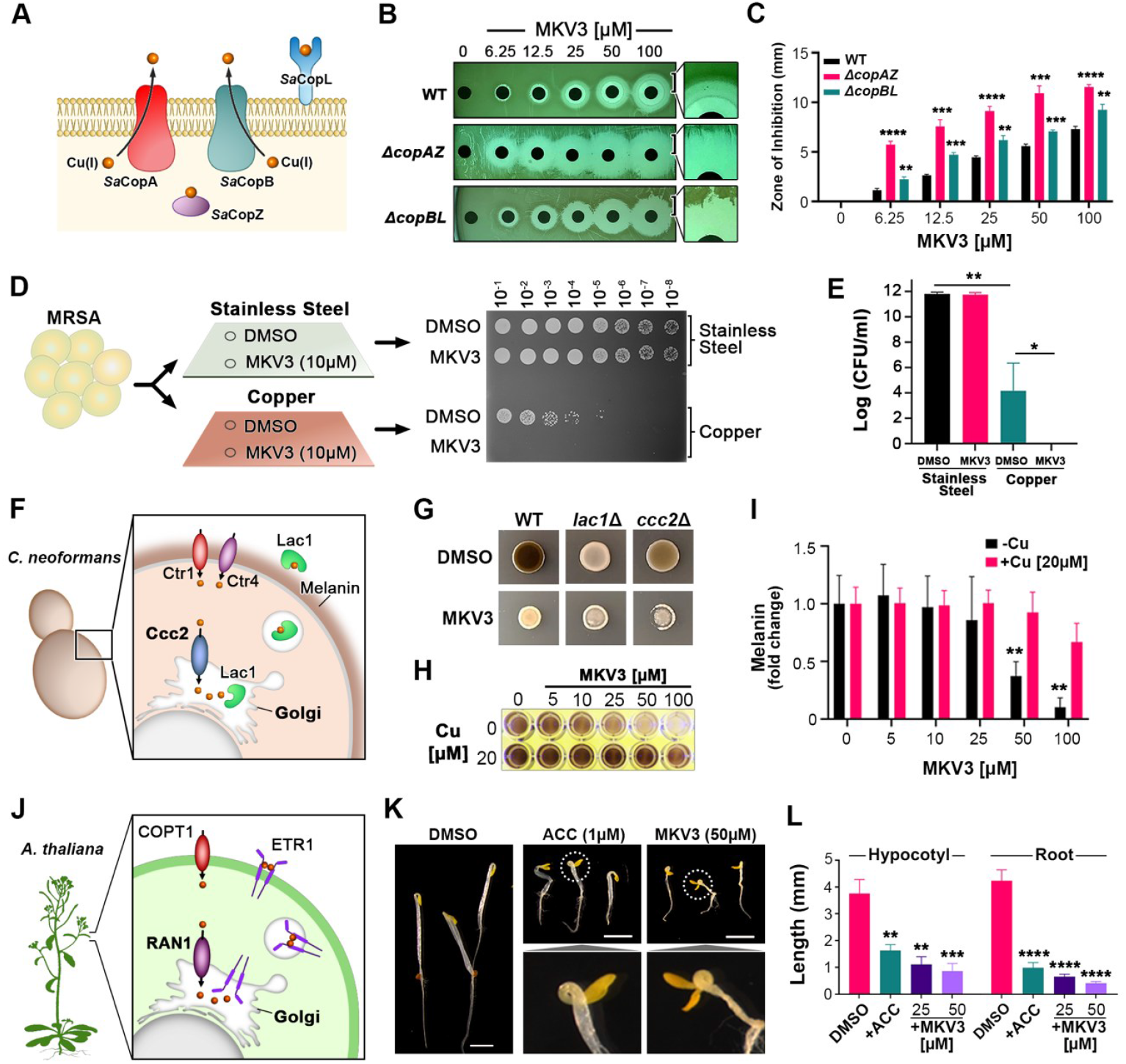
MKV3 targets Cu-ATPases across a broad evolutionary distance. (A) Schematic representation of the copper resistance system in *Staphylococcus aureus*, including *Sa*CopA, *Sa*CopB, *Sa*CopZ, and *Sa*CopL. (B) Zone-of-inhibition assay showing dose-dependent growth inhibition by MKV3 in *S. aureus* wild-type (WT), *ΔcopAZ*, and *ΔcopBL* strains plated on agar containing 1.5 mM CuCl_2_. (C) Quantification of the MKV3-dependent inhibition zones for WT, *ΔcopAZ*, and *ΔcopBL* strains. (D) Copper surface-mediated killing of MRSA is potentiated by MKV3. Wild-type MRSA in the presence of 10 µM MKV3 or DMSO control was spotted onto stainless steel or copper surfaces, incubated at room temperature for 10 minutes, and subsequently quantified by serial dilution plating. (E) Quantification of MRSA survival following MKV3 or DMSO treatment under the conditions shown in panel D. (F) Schematic illustration of the copper transport and melanin biosynthesis pathways in *Cryptococcus neoformans*, highlighting the roles of Lac1 and Ccc2. (G) Fungal pigmentation assessment on solid agar plates containing L-DOPA for wild-type (WT), *lac1*Δ, and *ccc2*Δ strains of *C. neoformans*, treated with MKV3 or DMSO and incubated at 30°C for 2 days. (H) Pigmentation assessment of *C. neoformans* WT strain in a 96-well assay treated with MKV3 alone or with 20 µM CuSO_4_, incubated at 30°C for 6 days. (I) Quantification of melanin production in *C. neoformans* WT strain treated with MKV3 in the presence or absence of 20 µM CuSO_4_. (J) Diagram of the ethylene signaling pathway in *Arabidopsis thaliana* highlighting RAN1 and ETR1. (K) Representative images of *A. thaliana* seedlings grown in medium containing DMSO, 1 μM 1-aminocyclopropane-1-carboxylic acid (ACC), or 50 μM MKV3. (L) Quantification of hypocotyl and root length in seedlings treated with DMSO, 1 μM ACC, 25 μM MKV3, or 50 μM MKV3. Data are presented as mean ± SEM. Statistical significance is indicated as follows: **p < 0.05; ***p < 0.001; ****p < 0.0001. Data represent the mean of three independent experiments.

We then assessed MKV3 activity in the human fungal pathogen *Cryptococcus neoformans*. The sole P_1B-1_-type ATPase in this organism, Ccc2, delivers copper into the Golgi complex to laccase 1 (Lac1), a secreted cuproenzyme that catalyzes melanin synthesis via the oxidative polymerization of catecholamines such as L-DOPA (53) (Fig. 6F). As expected, both the *ccc2*Δ and laccase mutant (*lac1*Δ) exhibited defective melanization compared to wild-type when grown on L-DOPA-supplemented agar (Fig. 6G). Consistent with inhibition of Ccc2, MKV3 blocked melanin production in wild-type *C. neoformans* in a dose-dependent manner (Fig. 6H and 6I).

Notably, MKV3 failed to block melanin production in media supplemented with excess copper, suggesting the inhibitor blocks copper delivery to Lac1 (Fig. 6H and 6I).

Next, we tested whether MKV3 inhibits the plant Cu^+^-ATPase RAN1 (HMA7) in *Arabidopsis thaliana*. RAN1 supplies copper to the ethylene receptor ETR1 during its maturation in the secretory pathway (Fig. 6J). Mutants of RAN1 germinated in the dark exhibit a characteristic “triple response” phenotype including an exaggerated apical hook and reduced hypocotyl and root elongation (22). This phenotype can be induced in wild-type seedlings by treatment with 1-aminocyclopropane-1-carboxylic acid (ACC), which increases ethylene production (54). As expected, ACC-treated seedlings displayed the triple response, and MKV3 treatment produced the same developmental defects, including a pronounced apical hook and shortened hypocotyls and roots (Fig. 6K and 6L). The ability of MKV3 to phenocopy the *ran1* mutant phenotype suggests MKV3 inhibits RAN1-mediated copper delivery to ETR1. Finally, we extended our analysis to *Trypanosoma cruzi*, the protozoan parasite responsible for Chagas disease (55). The genome of *T. cruzi* expresses a single Cu^+^-exporting ATPase, *Tc*CuATPase, whose expression increases upon copper exposure during the epimastigote stage (56). Treatment with MKV3 significantly increased the sensitivity of epimastigotes to copper, and exogenous copper significantly enhanced MKV3 toxicity (Fig. S11). Together, these data provide compelling evidence that MKV3 is a broad-spectrum inhibitor of Cu^+^-ATPases across bacteria, protozoa, fungi, plants, and animals, highlighting an evolutionarily conserved, druggable vulnerability within this essential class of P-type transporters.

## Discussion

Cu^+^-ATPases play central roles in regulating copper homeostasis in virtually all organisms, governing both biosynthetic metalation and the detoxification of excess cytosolic copper. Despite their fundamental biological importance and their documented roles in microbial pathogenesis, plant signaling, and human disease, no small-molecule inhibitors of Cu^+^-ATPases have previously been described. The present work addresses this gap by identifying MKV3 as the first small-molecule inhibitor of Cu^+^-ATPases, and by establishing that a conserved structural pocket can be exploited to achieve broad-spectrum inhibition across bacteria, fungi, plants, and animals.

Our *in-silico* screen for an inhibitor of Cu^+^-ATPases focused on the highly conserved Cu^+^ entry pocket at the cytoplasmic face of the membrane domain (Fig. 1C). This pocket is proposed to serve as a site of Cu^+^ delivery by cytosolic chaperones or N-terminal metal-binding domains, yet its potential as a druggable target had not previously been explored. Our finding of a competitive relationship between MKV3 and the N-terminal MBD5/6 domains of ATP7A (Fig. 2), supports a model in which the inhibitor occupies the same cavity used for physiological metal transfer to the CPC motif within the membrane translocation pathway. Cu^+^ delivery to the CPC motif is a known pre-requisite for coupled ATP hydrolysis (57) and consistent with our model, MKV3 blocked both the ATPase activity and Cu^+^ transport function of *Ec*CopA (Fig. 3A and 3E). Additionally, MKV3 decreased *Ec*CopZ-mediated Cu^+^ loading of *Ec*CopA in a manner consistent with selective blockade of transfer to the transmembrane CPC site while preserving Cu^+^ delivery to the N-terminal metal-binding domains (Fig. 3H). Taken together, these findings suggest a mechanism of inhibition in which MKV3 blocks Cu^+^ delivery to the catalytic CPC motif.

A distinguishing feature between MKV3 and the closely related but inactive analog MKV2 is the substitution of a thioamide sulfur with oxygen, suggesting that this single-atom change plays a critical role in inhibitor activity. Geometry optimization using *ab initio* quantum chemical calculations suggests MKV2 and MKV3 adopt distinct low-energy conformations (Fig. S12), consistent with the sulfur substitution altering the geometry and electrostatic properties of the scaffold that are more favorable for binding in the Cu^+^ entry pocket. Moreover, MKV3 binding to ATP7A was dependent on the Cu-coordinating cysteines of MBD5/6, consistent with the idea that the inhibitor preferentially engages a functional state of the pocket normally accessed during physiological Cu^+^ delivery. One speculative model is that the thioamide sulfur of MKV3 mimics aspects of sulfur-based interactions used by MBD5/6 during Cu^+^ transfer, without functioning as a Cu delivery vehicle itself.

The identification of a single P-domain residue as a determinant of MKV3 potency provided additional mechanistic insight and a rational foundation for future inhibitor design. Structural modeling predicted that this residue (glutamate in ATP7A and *Sa*CopA, lysine in ATP7B) forms an electrostatic interaction with a fluorine atom on MKV3 (Fig. 5). Substitution experiments confirmed that converting the glutamate to lysine in both human ATP7A and bacterial *Sa*CopA increased MKV3 affinity and potency, suggesting that a single charge-dependent contact modulates drug efficacy across species. These findings establish proof-of-principle that inhibitor potency can be tuned through targeted modifications and raise the possibility of engineering species-selective or pathogen-specific Cu^+^-ATPase inhibitors.

A major advance of this study is the demonstration that MKV3 produces phenotypes indistinguishable from genetic loss of Cu^+^-ATPase function across evolutionarily distant organisms. In mammalian cells, MKV3 blocked copper-stimulated trafficking of ATP7A, increased cellular copper burden, sensitized cells to copper toxicity, and triggered cuproptosis (Fig. 4A – 4H). These effects were accompanied by impaired ATP7A-dependent copper delivery to secretory pathway cuproenzymes, including lysyl oxidase and tyrosinase, both of which are critical for extracellular matrix remodeling and melanin biosynthesis, respectively (Fig. 4I – 4K). *In vivo*, MKV3 suppressed melanogenesis in a murine model of metastatic melanoma and zebrafish embryos, confirming target engagement in intact organisms (Fig. 4L – 4N).

The antimicrobial potential of MKV3 was demonstrated using the pathogenic USA300 strain of methicillin-resistant *S. aureus* where the compound inhibited growth in a copper-dependent manner and sensitized bacteria to the biocidal effects of solid copper surfaces (Fig. 6B – 6E). Notably, MKV3 sensitized USA300 in a manner consistent with inhibition of both the P_1B-1_-type Cu^+^-ATPase *Sa*CopA and the P_1B-3_-type *Sa*CopB, underscoring its ability to compromise multiple copper export systems within a single pathogen. Similarly, in the fungal pathogen *C. neoformans*, MKV3 blocked laccase 1-dependent melanin production, phenocopying the loss-of-function *ccc2*Δ mutant (Fig. 6G – 6I). In *A. thaliana*, MKV3 recapitulated the classic triple-response phenotype associated with constitutive ethylene signaling, consistent with inhibition of RAN1-mediated copper delivery to the ethylene receptor ETR1 (Fig. 6K and 6L). Collectively, these results establish that MKV3 targets a common mechanistic vulnerability shared across kingdoms and validate Cu^+^-ATPases as druggable targets in copper-dependent physiology.

The discovery that a single compound can inhibit Cu^+^-ATPases in both pathogenic microbes and mammalian cells opens new avenues for therapeutic intervention. In infectious disease, copper-based antimicrobial strategies have gained renewed interest as alternatives to conventional antibiotics, particularly for multidrug-resistant pathogens (58–60). Copper and its various alloys were the first solid surface products permitted by the US Environmental Protection Agency (EPA) to make public health antibacterial claims. MKV3’s ability to potentiate copper-mediated killing of MRSA offers a potential strategy to amplify the antimicrobial activity of solid copper surfaces used in healthcare and public settings to prevent nosocomial infections (61). Beyond antimicrobial applications, MKV3 may also have applications in oncology, where copper homeostasis is increasingly recognized as a determinant of tumor behavior and therapy response (3, 16). ATP7A and ATP7B have been linked to platinum drug resistance (20, 62–64), copper-dependent tumor growth (65), and the activity of secreted copper enzymes such as lysyl oxidase that contribute to extracellular matrix remodeling and metastasis (19, 66). Inhibition of these transporters by MKV3 is therefore expected to increase intracellular copper levels, impair copper delivery to key effectors, and potentially sensitize tumor cells to copper-dependent stress and chemotherapeutic agents.

In summary, the broad activity of MKV3 across bacteria, fungi, plants, and vertebrates underscores the evolutionary conservation in Cu^+^ ATPases, and opens the door to a new class of chemical tools and potential therapeutics aimed at modulating copper-dependent processes across biology. The evolutionary conservation of this pocket across bacteria, fungi, plants, and vertebrates establishes a general framework for the rational design of inhibitors with tunable potency and species selectivity. Together, these results position Cu^+^-ATPases as a new class of tractable drug targets and provide a foundation for chemical control of copper-dependent biology across kingdoms.

## Materials and Methods

Comprehensive details of all materials and experimental procedures employed in this study are provided in the SI Appendix, Materials and Methods. These encompass structure-guided docking, chemical synthesis, nuclear magnetic resonance (NMR), high-resolution mass spectrometry (HRMS), high-performance liquid chromatography (HPLC), microscale thermophoresis (MST), ATPase assays, proteoliposomal Cu(I) transport, inductively coupled plasma mass spectrometry (ICP–MS), cuproenzyme assays, immunoblotting, immunofluorescence, cell viability assays, zone-of-inhibition and copper contact-kill assays and other pertinent methodologies.

All animal experiments were conducted in accordance with institutional and national ethical guidelines and in compliance with protocols approved by institutional animal care and use committees (IACUC) of the University of Missouri and Texas A&M University.

## Acknowledgments

This work was supported by grants from the National Institutes of Health to M.J.P. (R01CA262664 and R01DK131190), V.M.G. (R35GM152102), and G.M. (R35GM128704). G.M. acknowledges support from the Robert A. Welch Foundation (AT-2073-20240404). CP acknowledges support by Duke University School of Medicine Bridge Funding (4532750) and Novo Nordisk Foundation (NNF21OC0070070). The authors thank Drs. M. Thomas Morgan and Christoph J. Fahrni (Georgia Institute of Technology) for providing the CTAP-3 probe. We thank Drs. Jeffrey Boyd (Rutgers University) and James Imay (University of Illinois Urbana-Champaign) for providing bacterial strains used in this study. ICP-MS measurements were performed in the OHSU Elemental Analysis Core (RRID: SCR_022746) with partial support from the NIH instrumentation grant S10RR025512. We also thank Dr. Jen Heemstra (Washington University in St. Louis) for generously providing access to the microscale thermophoresis (MST) instrument for binding experiments. The content herein is solely the responsibility of the authors and does not necessarily represent the official views of any funding agency.

