## Supplementary file for "A First-In-Class Broad Spectrum Inhibitor of Copper Exporting P_1B_-type ATPases"

### SI Appendix Materials and Methods

#### Structure preparation and molecular docking

The cryo-electron microscopy structure of frog *Xt*ATP7B (PBD 7SI3) from *Xenopus tropicalis* was used for structure preparation and molecular docking (1). The triad residues (D915, E778, M726) that are thought to facilitate copper entry into the membrane translocation pathway were selected to identify a druggable binding pocket. A grid of 22 Å × 22 Å × 22 Å centered on the triad residues was generated by the Receptor Grid Generation utility of the Glide module in the Schrödinger software suite (Schrödinger). A Maybridge Hitfinder library of $10^8$ compounds was used for docking onto the grid. All docking simulations were conducted using the Induced-Fit Docking (IFD) module of Schrödinger Suite (Schrödinger). The IFD used Glide (Schrödinger) and the refinement module in Prime (Schrödinger) to predict ligand binding modes in the binding pocket of the protein. The IFD optimized the side chain conformation to best determine the docking poses. The pose with the best IFD score was selected for comparison purposes.

#### Chemical synthesis

*General procedures*

All commercially available reagents and solvents were used as is without further purification. All reactions unless otherwise stated were run under inert argon atmosphere. Flash chromatography was performed either manually or on a C-805 Buchi chromatography system. NMR spectra were obtained on a Bruker 600 MHz or a 400 MHz spectrometer. High resolution mass spectra (HRMS) and analytical HPLC traces were obtained from an Agilent 6230 LC/TOF mass spectrometer with an ESI source coupled with an Agilent Infinity 1260 HPLC system running in reverse phase on a Poroshell 120 EC-C18 (4.6 × 50 mm, 2.7 µm) using water with 0.1% formic acid as solvent A and acetonitrile with 0.1% formic acid as solvent B. For HRMS, the gradient for elution varied from 30% B to 95% B over 7 minutes at a flow rate of 0.5 mL/min. For analytical HPLC, the gradient for elution varied from 30% B to 95% B over 8 minutes and kept at this gradient for 4 minutes at a flow rate of 0.6 mL/min for a 12 min run.

*Synthesis of Intermediate 1a*

Cyanoacetic acid (500 mg, 5.87 mmol) and 3-trifluoromethyl aniline (631 mg, 3.92 mmol) was added to 10 mL of dichloromethane and to this solution was added triethylamine (1.63 mL, 11.76 mmol) and the solution was cooled in an ice bath, followed by the addition of 50 wt % of propylphosphonic anhydride in ethyl acetate (3.74 mL, 5.87 mmol) and the reaction was stirred overnight. After TLC showed the complete consumption of 3-trifluoromethyl aniline, the mixture was subjected to aqueous workup followed by extraction with ethyl acetate. The organic layers were evaporated, and the crude solid was subjected to column chromatography with hexanes/ethyl acetate to give 840 mg of intermediate 1a as a white solid. Yield: 94 %; $^1$H NMR (600 MHz, DMSO) δ 10.63 (s, 1H), 8.02 (s, 1H), 7.73 (d, *J* = 8.0 Hz, 1H), 7.58 (t, *J* = 8.0 Hz, 1H), 7.45 (d, *J* = 7.8 Hz, 1H), 3.95 (s, 2H); $^{13}$C NMR (151 MHz, DMSO) δ 161.76, 139.12, 130.24, 129.59 (q, *J* = 31.7 Hz), 124.00 (q, *J* = 272.3 Hz) ,122.84, 120.27(m, *J* = 3.7 Hz), 115.30 (m, *J* = 4.0 Hz), 26.89; HRMS (ESI) $[M+H]^+$Calculated for $C_{10}H_8F_3N_2O^+$ is 229.0583, found 229.0592.

*Synthesis of MKV2*

100 mg (0.44 mmol) of amide 1a was dissolved in 1 ml of DMF in a one-dram vial and to this was added 32 mg (0.57 mmol) of KOH was added and after dissolving the KOH the reaction mixture was sealed under argon with a cap containing a septum. The reaction mixture was cooled in ice and a solution of 63 µl (0.53 mmol) of 2-chlorophenyl isocyanate in 0.5 ml of DMF was added dropwise and the reaction was stirred and allowed to warm up to room temperature. The reaction mixture was stirred at room temperature for 40 hrs. After the starting material was consumed the reaction was quenched by pouring it into a mixture of ice and 1N HCl. The crude product was obtained as a light-yellow solid. The crude product was recrystallized from a mixture of DCM and Hexanes to give 138 mg of MKV2 as a white solid. MKV2 was obtained as a ~1:1 mixture of keto and enol tautomers. Yield : 82.6 %; $^1$H NMR (600 MHz, $CD_3CN$) δ 9.21 (s, 1H), 8.96 (s, 1H), 8.32 (s, 1H), 8.11 (s, 1H), 7.97 (s, 2H), 7.82 (s, 1H), 7.80 – 7.67 (m, 3H), 7.53 (m, 6H), 7.38 (dd, *J* = 17.2, 9.3 Hz, 2H), 7.29 (t, *J* = 7.6 Hz, 1H), 7.23 (s, 1H), 5.02 (s, 1H); $^{13}$C NMR (151 MHz, $CD_3CN$) δ 173.06, 172.64, 160.67, 160.04, 139.09, 137.94, 134.69, 133.78, 131.09, 130.84, 130.67, 130.61, 129.12, 128.82, 128.78, 127.95, 127.53, 127.36, 126.74, 125.97, 125.21, 124.92, 124.62, 124.17, 122.90, 122.65, 120.44, 116.73, 114.68, 58.90, 48.92; $^{19}$F{$^1$H} NMR (376 MHz, $CD_3CN$) δ -63.30, -63.35; HRMS (ESI) m/z: $[M+H]^+$Calculated for $C_{17}H_{12}ClF_3N_3O_2$ 382.0565; found 382.0563.

*Synthesis of MKV3*

Amide 1a (200 mg, 0.88 mmol) was dissolved in 1.5 ml of DMF in a one-dram septa sealed vial and to this KOH (64 mg,1.14 mmol) was added. The reaction mixture was sealed under argon and stirred until most of the KOH dissolved. The reaction mixture was cooled in ice and to this was added dropwise via a syringe, a solution of the 2-chlorophenyl isothiocyanate (178 mg, 1.05 mmol) dissolved in 0.5 ml of DMF and the reaction mixture was allowed to warm up to room temperature. The reaction mixture was stirred at room temperature for 48 hr. After TLC showed the complete consumption of amide 1a, the reaction was quenched by pouring it into ice cold dilute HCl. The product precipitated out as a pale-yellow solid. The crude product was further purified by column chromatography using hexanes and ethyl acetate gradient elution. The pure fractions were collected and subjected to an aqueous HCl wash. The organic layer was evaporated to obtain 220 mg MKV3 of a yellow solid. Yield: 63.2%; HPLC purity : 98%; $^{1}$H NMR (600 MHz, 10:1 DMSO-$d_6$:$D_2O$) δ 8.31 (s, 1H), 7.96 (s, 1H), 7.70 (d, *J* = 7.5 Hz, 1H), 7.46 (t, *J* = 8.0 Hz, 1H), 7.42 (d, *J* = 7.9 Hz, 1H), 7.28 (d, *J* = 7.6 Hz, 1H), 7.23 (t, *J* = 7.6 Hz, 1H), 7.08 (t, *J* = 7.2 Hz, 1H); $^{13}$C NMR (151 MHz, 10:1 DMSO-$d_6$:$D_2O$) δ 187.98, 167.29, 140.89, 138.12, 130.25, 129.98, 129.60, 127.63, 126.85, 125.91, 125.76, 124.72, 124.19, 123.95, 119.12, 116.58, 78.13; $^{19}$F{$^{1}$H} NMR (376 MHz, 10:1 DMSO-$d_6$:$D_2O$) δ -61.16 HRMS (ESI) $[M+H]^+$Calculated for $C_{17}H_{12}ClF_3N_3OS^+$ is 398.0336 ; found 398.0337.

**Microscale Thermophoresis (MST) assays**

MST was used to measure compound binding affinities to human ATP7A/B and mutant variants that were GFP-tagged at the N-terminus. HEK-293 cells were lysed by sonication in PBS containing EDTA-free protease inhibitors. Lysates were clarified by centrifugation at 500 × g for 15 min, protein concentrations were quantified using a Bradford assay kit (BioRad), and equal amounts of protein were incubated with MKV3. The resulting reaction mixtures were loaded into capillaries, and thermophoresis was monitored at 20% LED power, high MST power with 20 s MST-on time. Data analysis was performed using MO. Affinity software (version 2.3) (NanoTemper Technologies, CA) and plotted using Prism (Version 6.0) (GraphPad Inc., La Jolla, CA).

**Expression and purification of *Ec*CopA *and Ec*ZntA**

Recombinant *Ec*CopA was expressed and purified from *E.coli* as previously described with minor modifications (2). Briefly, a pET-52b(+) expression vector (Novagen) encoding for wild-type *Ec*CopA (UniProt accession number: Q59385) featuring an N-terminal STREP-tag II (Genscript Inc.) was transformed into BL21-Gold(DE3) competent cells (Agilent Technologies). Cells were inoculated in Terrific Broth (TB) media supplemented with 1% glycerol (v/v) in the presence of ampicillin (50 μg/mL) and grown at 37°C overnight with orbital shaking (140 rpm). The overnight preculture was inoculated at 6% (v/v) in fresh TB media and cells grown at 37 °C under constant shaking (150 rpm) until they reached $OD_{600}$ = 2. Cell cultures were subsequently cooled to 25 °C and protein expression was induced by addition of isopropyl β-D-1-thiogalactopyranocide (IPTG) to a final concentration 0.3 mM. The protein was expressed for 18 h at 25 °C, under agitation. Cells were harvested by centrifugation (20 min, 4 °C, 14000 x*g*; Thermo Scientific Sorvall LYNX 6000 centrifuge) and the cell pellet was resuspended in lysis buffer (20 mM Tris/HCl pH 8, 150 mM NaCl, 5 mM $MgCl_2$, 30 μg $mL^{-1}$ deoxyribonuclease I from bovine pancreas (Sigma-Aldrich), and 2x EDTA-free protease inhibitor cocktail tablets (Thermo Scientific). Cells were lysed in an ice-chilled microfluidizer at 20000 psi by running the cell suspension through a Z-shaped diamond chamber (Microfluidics M-110P). Cell debris was removed by centrifugation (20 min, 4 °C, 20000 x *g*; Thermo Scientific Sorvall LYNX 6000 centrifuge). The membrane fraction containing *Ec*CopA was isolated by ultracentrifugation (1 h, 4 °C, 205100 x*g*, Beckman Optima XPN80). The membrane pellet was resuspended in buffer (20 mM Tris/HCl pH 8, 500 mM NaCl, 1% (w/v) glycerol, 1x EDTA-free protease inhibitor cocktail) to a final concentration of 1 g of cells per mL of buffer. Membrane suspensions were flash-frozen in liquid nitrogen and stored at −80 °C until purification. For purification, a 5 mL membrane suspension aliquot was diluted to a final volume of 50 mL with ice-cooled protein extraction buffer (20 mM Tris/HCl pH 8, 500 mM NaCl, 1 mM EDTA, 5 mM β-mercaptoethanol, 1% (w/v) n-dodecyl-β-D-maltoside (DDM; Anatrace) and 1x EDTA-free protease inhibitor cocktail), and vigorously stirred for 1 h at 4 °C to extract membrane proteins. Residual membrane debris was removed by ultracentrifugation (20 min, 4 °C, 205100 x*g*). The supernatant containing detergent-solubilized proteins was loaded onto a 5 mL StrepTrap column (Cytiva) equilibrated with wash buffer (20 mM Tris/HCl pH 8, 500 mM NaCl, 1 mM

EDTA, pH 8, 1 mM dithiothreitol (DTT), 0.05% (w/v) DDM) using an AKTA Pure FPLC system (Cytiva). Unbound proteins were removed with 20 CV of wash buffer. *Ec*CopA bound to the StrepTrap resin was eluted in 6 CV (30 ml) of buffer containing D-desthiobiotin (20 mM Tris/HCl pH 8, 500 mM NaCl, 1 mM EDTA, pH 8, 1 mM dithiothreitol (DTT), 0.05% (w/v) DDM, and 2.5 mM D-desthiobiotin). The eluted protein was immediately injected into a pre-equilibrated Superdex 200 10/300 column (Cytiva) using sizing buffer (20 mM MOPS/NaOH pH 7, 500 mM NaCl, 1 mM ascorbic acid or 1mM DTT, 0.05% (w/v) DDM). Protein purity was verified by SDS-PAGE (4–15% Tris-Glycine Mini-PROTEAN gels, BioRad). Protein concentration was determined by absorption at 280 nm ($\varepsilon_{280}$= 70275 $M^{-1}$ $cm^{-1}$) on a Nanodrop spectrophotometer (Thermo Scientific Nanodrop one). The final purified protein was stored at −80 °C after flash-freezing with $LN_2$, until further use.

*EcZnt*A was expressed and purified from *E.coli* using a pBAD vector (Thermo Fisher) encoding for *wild-type Ec*ZntA (UniProt accession number: P37617) featuring a C-terminal $His_6$tag. Protein expression was conducted in TB media without glycerol for 20 hours at 23°C, with the exception that the induction was performed by adding 0.02 % (w/v) arabinose. For purification, a 10 mL membrane suspension aliquot was diluted to a final volume of 50 mL with ice-cold protein extraction buffer (20 mM Tris/HCl pH 8, 500 mM NaCl, 25 mM imidazole, 5 mM β-mercaptoethanol, 1% (w/v) n-dodecyl-β-D-maltoside (DDM; Anatrace) and 1x EDTA-free protease inhibitor cocktail (Thermo Scientific), and vigorously stirred for 1 h at 4 °C to extract membrane proteins. Residual membrane debris and insolubilized membranes were removed by ultracentrifugation (20 min, 4 °C, 205100 x*g*).The supernatant containing detergent-solubilized proteins was loaded (0.5 mL $min^{-1}$ maximum speed) onto two 5 mL HisTrap FF affinity columns (Cytiva) pre-equilibrated with wash buffer (20 mM Tris/HCl pH 8, 500 mM NaCl, 35 mM imidazole, 1 mM dithiothreitol (DTT), 0.05% (w/v) DDM), using an AKTA Pure FPLC system (Cytiva). The columns were washed with 35 CV wash buffer to remove unbound and loosely bound protein impurities. *Ec*ZntA bound to the Ni-NTA resin was eluted with a linear imidazole gradient (0–100%) obtained by mixing wash and elution buffer (20 mM Tris/HCl, pH 8, 500 mM NaCl, 500 mM imidazole, 0.05% (w/v) DDM, 1 mM DTT) over 8 CV. The eluted protein buffer was immediately exchanged to remove imidazole by loading samples onto an equilibrated HiPrep 26/10 desalting column (Cytiva) and eluted with desalting buffer (20 mM MOPS/NaOH pH 7, 500 mM NaCl, 1 mM DTT, 0.05% (w/v) DDM). The eluted protein was concentrated on 100,000 MWCO filter concentrators (Sartorious VIVASPIN 20) by centrifugation (4000 ×*g*, 4 °C, Thermo Scientific) to 500 µL (~ 2-3 mg/mL). *Ec*ZntA was further purified by size exclusion chromatography to remove any low molecular weight impurities or aggregated protein, by injecting the protein sample into an equilibrated Superdex 200 10/300 column (Cytiva) and eluted in sizing buffer (20 mM MOPS/NaOH pH 7, 500 mM NaCl, 1 mM DTT, 0.05% (w/v) DDM). Protein purity was verified by SDS-PAGE (4–15% Tris-Glycine Mini-PROTEAN gels, BioRad). Protein concentration was determined by absorption at 280 nm ($\varepsilon_{280}$= 41940 $M^{-1}$ $cm^{-1}$) on a Nanodrop spectrophotometer (Thermo Scientific Nanodrop one).

**Determination of *Ec*CopA and *Ec*ZntA specific ATPase activity in the presence of MKV3 and MKV2.** All solutions were prepared in Chelex 100-treated milliQ water. Wild type *Ec*CopA (34 µL,~0.5 mg/mL) was placed in a 96-well plate and mixed with 10 mM $MgCl_2$, 2 mM cysteine and10 µM $CuCl_2$. Prior to adding 1 mM ATP to start reaction, the wells containing the reaction mixtures were treated with different concentrations of inhibitor in DMSO (0-5000 nM final concentration). The reaction mixtures were incubated at 37 °C for 20 minutes, with shaking (300 rpm, Eppendorf Thermomixer). The inorganic phosphate ($P_i$) released during the reaction was quantified using a Malachite green phosphate assay kit (Sigma Aldrich, MAK307). After incubating the reaction mixtures with the Malachite green reagent for 5-10 minutes at room temperature, the generated malachite green-phosphate complex was quantified by determining the absorbance at 620 nm using a Tecan Spark 20 M plate reader. Control experiments were conducted in the absence of $MgCl_2$ (an essential cofactor for ATP hydrolysis) and Cu. The released inorganic phosphate was calculated using a standard $P_i$ calibration curve and data presented as specific ATPase activity (nmol of $P_i$ released per minute per 1 mg of protein). Similar experiments were conducted for *Ec*ZntA, with the exception that reaction mixtures were supplemented with 10 µM $ZnCl_2$ instead of 10 µM $CuCl_2$. As maintaining a strong reducing environment with Zn(II) is not necessary, cysteine (supplemented to reduce Cu(I) to Cu(I)) was not included in the ATPase assays for *Ec*ZntA.

**Real-time determination of Cu(I) transport in EcCopA proteoliposomes in the presence of MKV3 and MKV2**

*Ec*CopA proteoliposomes were prepared as described previously (2). Proteoliposomes and control liposomes were diluted to a final lipid concentration of 12.5 mg $mL^{-1}$ in transport assay buffer containing 20 mM MOPS/NaOH pH 7.0, 100 mM NaCl, 1 mM ascorbic acid. The Cu(I)-selective fluorescence detector probe CTAP-3 (10 mM in ultra-pure water) was added to final concentration of 20 µM. CTAP-3 encapsulation in the liposome lumen was obtained by 3 freeze–thaw cycles and subsequent proteoliposomes/control liposomes extrusion through 1 µm, 0.4 µm and 0.2 µm polycarbonate (PC) filter membranes, using 1mL gas-tight syringes connected to a mini-extruder (Avanti Polar Lipids). The CTAP-3 containing liposome pellets were collected by ultracentrifugation (160000 x*g*, 45 min, 4°C; Sorvall Mx 120+ micro-ultracentrifuge) and the supernatant containing excess CTAP-3 probe was removed. Proteoliposomes were washed three times using 1mL of the transport assay buffer and collected by additional ultra-centrifugation/resuspension cycles to remove all the unencapsulated probe, and finally resuspended in transport assay buffer at a final *Ec*CopA concentration of 0.5 mg $mL^{-1}$. All Cu(I) containing solutions used for transport assays were made oxygen-free on a Schlenk-line by three vacuum/nitrogen cycles. Cu(I) stocks were freshly prepared prior to each experiment by diluting a 100 mM Cu(I) stock obtained by solubilizing $Cu(CH_3CN)_4]PF_6$ in 100% acetonitrile, in an anaerobic glove box purged with constant $N_2$ gas flow. The Cu(I) transport assays were conducted at 37°C in a sub-micro quartz cell (Starna Cells) using a Fluoromax-4 spectrofluorometer (Horiba Scientific). Proteoliposomes and control liposomes aliquots (127.4 µL) were supplemented with 1 mM ATP (100 mM stock) and 10 mM $MgCl_2$ (1 M stock). The reaction was initiated by the addition of 1.3 µL of incubated Cu(I) stock (final concentration: 20 µM). The time-dependent fluorescence change was measured for 250 s in 0.1 s intervals ($\lambda_{ex}$ = 365 nm, slit width = 2 nm; $\lambda_{em}$ = 450 nm, slit width = 2 nm). $(F - F_0)/ F_0$ was calculated using the fluorescence before addition of Cu(I) as $F_0$. Control experiments were performed without ATP. To assess the effects of MKV3 and MKV2 on real-time Cu(I) translocation by *Ec*CopA in proteoliposomes, the transport assays were repeated in the presence of 5 µM MKV3 or MKV2.

**Effect of MKV3 on *Ec*CopZ-mediated Cu(I) transfer to *Ec*CopA**

The effect of MKV3 on *Ec*CopA Cu(I) binding stoichiometry was determined by Inductively Coupled Plasma Mass Spectrometry (ICP-MS) analysis. All the steps, except for the final SEC on a Superdex 75 10/300 GL column, were performed in an anaerobic glove box purged with constant $N_2$ gas flow. All solutions were made oxygen-free on a Schlenk-line by three vacuum/nitrogen cycles. DTT present in the purified *Ec*CopA samples was removed prior to Cu(I) binding experiments by injecting concentrated protein stocks (40 µM, 1.5 mL) on a 5 mL HiTrap desalting column and eluted in assay buffer (20 mM MOPS pH=7, 500 mM NaCl, 0.05% (w/v) DDM) in the glovebox. DTT-free *Ec*CopA (35 µM, 250 µL) was supplemented with one equivalent (mol:mol) of MKV3 (from a 100 mM stock in DMSO) or DMSO as control. In parallel, purified recombinant untagged *Ec*CopZ (105 µM, 250 µL) in 20 mM MOPS pH=7, 500 mM NaCl, was loaded with one molar equivalent of Cu(I) (100 mM stock in $CH_3CN$) to form the EcCopZ-Cu(I) complex. *Ec*CopZ-Cu(I) (250 µL) was added at 3:1 (mol:mol) molar ratios to *Ec*CopA containing MKV3 (250 µL) and incubated at room temperature for 10 min. The protein mixtures were subsequently injected into a pre-equilibrated Superdex 75 10/300 GL column (Cytiva) to separate Cu(I)-bound *Ec*CopA from free *Ec*CopZ or Cu(I). *E*cCopA was collected at the column void volume. *Ec*CopA concentration was determined via Bradford assays and copper content determined upon protein digestion in high-purity $HNO_3$ (8% (w/v); Sigma-Aldrich) at 85 °C for 12 h. The $HNO_3$ concentration was subsequently adjusted to 3% (w/v) by dilution with Chelex 100-treated MilliQ water. The copper was quantified on an Agilent 7900 ICP mass spectrometer connected to a CETACASX-500 auto-sampler for sample injection. Parallel experiments were performed by incubating DTT-free EcCopA (35 µM, 250 µL) treated with one molar equivalent of MKV-3 inhibitor or DMSO, with either *Ec*CopZ-Cu(I) (35 µM) at 1:1 molar ratio, or “free” Cu(I) (105 µM; molar ratio of 3:1 *vs. Ec*CopA) in *Ec*CopZ buffer (250 µL).

**Mammalian Cell Culture and plasmid transfections**

The 4T1 and B16-F10 cells were sourced from the American Type Culture Collection and cultured in complete medium consisting of Dulbecco's Modified Eagle Medium (DMEM; Life Technologies) supplemented with 10% (v/v) FBS, 2 mM glutamine, and 100 U/mL penicillin-streptomycin (Life Technologies) in a 5% $CO_2$ atmosphere at 37 °C. *Atp7a* knockout B16-F10 cells were generated as previously described (3). The GFP-tagged human ATP7B expression plasmid was provided by Dr.

Roman Polischuk (4). All GFP-tagged human ATP7A constructs and mutant variants were synthesized by Gene Universal and subcloned into pcDNA3.1. All plasmids were verified by DNA sequencing.

**Cellular copper measurements**

Cellular copper measurements were determined as previously described (5). Briefly, cell pellets were digested in 15 ml metal-free centrifuge tubes (VWR 89049-170) with 100 µl of concentrated $HNO_3$ (trace metal grade, Fisher). Samples were heated to 90°C for 45 mina and subsequently allowed to cool and finish digesting at room temperature overnight before 1% HNO3 (trace metal grade, Thermo Fisher Scientific) was added to a total volume of 2 ml and measured undiluted. Inductively coupled plasma mass spectroscopy (ICP-MS) analysis was performed using an Agilent 7700x equipped with an ASX 500 autosampler. The system was operated at a radio frequency power of 1550 W, an argon plasma gas flow rate of 15 L/min, Ar carrier gas flow rate of 0.9 L/min. Elements were measured in kinetic energy discrimination (KED) mode using He gas (4.3 ml/min). Data were quantified using a 11-point calibration curve made from serial dilutions of a multi-element standard (VHG-SM70B-100). For each sample, data were acquired in triplicates and averaged. A coefficient of variance (CoV) was determined from frequent measurements of a sample containing ~10 ppb. An internal standard (Sc, Ge, Bi) continuously introduced with the sample was used to correct for detector fluctuations and to monitor plasma stability. Accuracy of the calibration curve was assessed by measuring NIST reference material (water, SRM 1643e).

**Tyrosinase activity assays**

*In-situ* tyrosinase activity in B16-F10 cells was measured as previously described (6). For quantitative determination of tyrosinase activity, cells grown in 96-well trays were washed with ice-cold PBS and lysed in 30µL RIPA buffer with protease inhibitors (Thermo Scientific, catalog# A32955) at 4°C for 30 minutes. A 70 µL volume of reaction buffer containing 50 mM phosphate buffer (pH 6.8) and 0.05% L-3,4-dihydroxyphenylalanine (L-DOPA) was added to wells and samples were incubated at 37°C for 60 minutes. Dopachrome formation was measured spectrophotometrically at 475 nm.

**Lysyl Oxidase activity assay**

Lysyl oxidase (LOX) activity was measured in the conditioned media of 4T1 breast cancer cells as previously described (7) using a kit from Abcam (ab112139) according to the manufacturer's instructions. Cells were cultured in phenol-red free DMEM supplemented with 10% FBS, 2.5% penicillin-streptomycin, non-essential amino acids, and L-Glutamine. Indicated concentrations of MKV3 or vehicle (1% DMSO) were replenished with daily media changes over 5 days.

**Immunoblot assays**

Mammalian and bacterial cell lysates were prepared by sonicating cell pellets in their respective lysis buffers: RIPA buffer supplemented with protease inhibitor cocktail (Thermo Scientific, catalog# A32955) for mammalian lysates, and 1X lithium dodecyl sulfate (LDS) buffer for bacterial lysates. For bacterial lysates, the bacterial pellet was collected when the bacterial growth $OD_{600}$ reached ~0.37 and was pelleted at this point, followed by addition of 1X LDS buffer. Fifty micrograms of each lysate were fractionated on 12% SDS-PAGE and transferred onto nitrocellulose membranes. After blocking with 5% non-fat milk in TBST (Tris-buffered saline with 0.1% Tween 20), membranes were incubated at 4°C overnight with primary antibodies. ATP7A antibodies used for Western analysis were raised against the last 33 amino acids of the C-terminus of the ATP7A protein (Bethyl Laboratories) as described in (8), Anti-GFP (D5.1) Rabbit mAb (Cell Signaling #2956), Anti-DLAT recombinant monoclonal antibody (clone ARC1314; Thermo Fisher Scientific #MA5-35723), Anti-β-Tubulin I Mouse mAB (Millipore Sigma #T7816) for mammalian lysates, and StrepTag™ II Monoclonal Antibody mAb (Millipore Sigma #71590-M) and Anti-GAPDH Mouse mAB (Abcam #ab125247) for bacterial lysates. For DLAT immunoblotting, lysates were not boiled and no reducing agents were added to the sample buffer to prevent disruption of DLAT oligomers by heat or reduction of disulfide bonds. Anti-mouse (#7076) or anti-rabbit (#7074) horseradish peroxidase-conjugated secondary antibodies (Cell Signaling) were used accordingly. Blots were developed using the SuperSignal West Pico Substrate according to the manufacturer's instructions (Pierce).

**Immunofluorescence microscopy**
4T1 cells on sterile glass coverslips were pre-treated for 16 hours at 37°C in DMEM media containing 10 µM of the copper chelator bathocuproinedisulfonate (BCS). The following day, cells were treated for 4 hours with either 2 µM Cu ± 10 µM MKV3, or 1% DMSO vehicle control. Cells were then washed twice with 1 ml of ice-cold PBS and fixed for 10 minutes at room temperature using 4% paraformaldehyde in PBS. Cells were then permeabilized with 0.1% Triton X-100 in PBS for 8 minutes and then blocked for 1 hour with 1% BSA in PBS. Next, cells were probed with an anti-ATP7A antibody, as described previously (8). Following three washes in PBS, cells were incubated with Alexa 488-conjugated anti-rabbit IgG (Invitrogen, catalog# A-11034) as secondary antibodies. Subsequently, cells were stained with 4′,6-diamidino-2-phenylindole (DAPI) to visualize nuclei, washed three times with PBS, and mounted. Imaging was performed using a Leica SP8 MP spectral scanning confocal microscope.

**Mammalian cell viability assays**
Cells were seeded into 12-well trays (30,000 cells/well) into basal media containing either MKV3 and/or copper at indicated concentrations. Cell survival was measured using a Crystal Violet assay, as described previously (9).

***E. coli* LEM33 strain complementation and MKV3 kill assays**
Expression constructs for wildtype and E481K variants of *Sa*CopA (Uniprot accession: Q2FV64) were synthesized commercially (Gene Universal) and subcloned into a pET-52b(+) expression vector (Novagen) and transformed into electrocompetent LEM33 *E.coli* for complementation analysis. Overnight cultures of LEM33 complemented strains in Luria broth were diluted 1:100 and grown at 37°C with agitation at 250 RPM until reaching an $OD_{600}$ of ~0.6. Isopropyl β-D-1-thiogalactopyranoside (IPTG) was added to a final concentration of 0.3 mM, and the cultures were further incubated overnight at 18°C with agitation at 250 RPM. For liquid broth experiments, LEM transformants were inoculated to a starting $OD_{600}$ of approximately ~0.037 under respective treatment conditions and incubated overnight at 37°C with agitation at 250 RPM. For plate assays, 1:10 serial dilutions of bacterial cultures were prepared and 10 µl of each dilution was spotted onto plates.

**Bacterial zone inhibition assays**
Methicillin-resistant *S. aureus* (MRSA) strain (USA300), *ΔcopBL* and *ΔcopAZ* were cultured in nutrient broth 1 (NB1) medium to an optical density ($OD_{600}$) of ~0.4, diluted 1:100 and plated onto NB1-agar ± 1.5 mM copper. After a 15-minute incubation at 37°C, sterile filter paper discs (Whatman, #1) soaked with either 10 µL of the indicated MKV3 concentration or DMSO vehicle were applied to the plates. Plates were further incubated for 16 hours at 37°C. The radius of the inhibited circular zone was measured and plotted for each treatment.

**Copper contact kill assays**
Methicillin-resistant *S. aureus* (MRSA) strains were cultured overnight in NB1 media, centrifuged at 2500 x g for 5 minutes, washed once with 10 ml of 100 mM Tris-Cl buffer (pH 7), and resuspended in 3 ml of the same buffer to obtain an $OD_{600}$ of ~0.3. Indicated concentrations of MKV3 or DMSO vehicle were mixed with the bacteria and 50 µl of the bacterial mixture was applied to either a copper surface (99.9%) or stainless-steel surface (K&S precision metals) at room temperature. At indicated times, ten-fold serial dilutions were prepared and 10 µL of each dilution was either spotted onto NB1 agar plates or added to NB1 medium and incubated for 16 hours at 37°C. Plate images were captured using the iBright 1500 imaging system. NB1 cultures were quantified ($OD_{600}$) and graphically represented.

**Determining $CC_{50}$ of MKV3 and $CuSO_4$ on *T. cruzi* Dm28c Epimastigotes**
The experiments were conducted using the *Trypanosoma cruzi* Dm28c strain. *T. cruzi* epimastigotes were routinely maintained in mid-log phase by periodic dilutions in Liver Infusion Tryptose (LIT) medium consisting of 5 g/L Liver Infusion Broth BD DIFCO™ # 226920, 5 g/L GranuCult® prime Tryptose Broth Millipore #1.03866, 68 mM NaCl, 5.3 mM KCl, 22 mM $Na_2HPO_4$, and 0.4 % (w/v) glucose, adjusted to pH 7.4. This medium was supplemented with 10% FBS (Internegocios S.A. - Buenos Aires, Argentina) and 5 µM hemin (LIT-10 % FBS-5 µM hemin medium) at 28°C. $EC_{50}$ determinations were performed in MW96 culture plates with a final volume of 100 µL per well. Parasites were seeded at a density of $2 \times 10^6$ epimastigotes/mL in LIT medium supplemented with MKV3 (0-80 µM) and $CuSO_4$ (0-100 µM) and

incubated for 72 hours. Cell growth was determined by cell counting using an automatized hemocytometer (WL 19 Counter AA, Weiner Lab, Rosario, Argentina) adapted to count epimastigotes at the White Blood Cell channel.

**Cryptococcus neoformans melanization plate assay**

For assessment of melanization on plates, cells were cultured and spotted on to L-3,4-dihydroxyphenylalanine (L-DOPA) containing plates as previously described in (10). To measure effects of MKV3 on melanization, the L-DOPA medium was supplemented with 25 µM MKV3 or DMSO. L-DOPA plates were incubated at 30°C for 2 days.

**Cryptococcus neoformans 96 well melanization assay**

For assessment of melanization in liquid culture, overnight cultures in yeast peptone dextrose media (YPD) were washed once with L-DOPA containing medium (7.6 mM L-asparagine monohydrate, 5.6 mM glucose, 22 mM $KH_2PO_4$, 1 mM $MgSO_4$X $7H_2O$, 0.5 mM L-DOPA, 0.3 µM thiamine-HCl, 20 nM biotin, pH 5.6). Cells were resuspended in L-DOPA medium and set to an $OD_{600}$ of 0.4. To avoid effects of DMSO on melanization, a 125x concentrated MKV3 inhibitor stock dilution series was made. 248 µL of diluted cells were aliquoted in a 96 well plate and mixed with 2 µL of 125x concentrated MKV3 dilutions (in DMSO, final concentrations in well: 0, 5 µM, 10 µM, 25 µM, 50 µM, 100 µM). For $CuSO_4$ supplementation, cells were cultured and washed as described above and then resuspended in L-DOPA medium to an $OD_{600}$ of 0.8 (=2x concentrated cells). In a 96 well plate 124 µL of 2x cells were mixed with 2x concentrated $CuSO_4$ solution (in L-DOPA medium, final concentrations in well 0 µM, 20 µM, 50 µM). Then, wells were supplemented with 2 µL of 125x concentrated MKV3 dilutions (in DMSO, final concentrations: 0, 5 µM, 10 µM, 25 µM, 50 µM, 100 µM). Plates were sealed using a Breathe-Easy plate membrane (Diversified Biotech) and incubated for 6-7 days at 30°C. Plates were imaged with the Azure 600 imager (Azure Biosystems) using the translucid sample setting. For further analysis, the images were converted to 513 nm greyscale images via the Azure Image software. The mean grey scale value was measured for each well using the ImageJ/Fiji software. To calculate the MKV3 inhibitory concentration, the mean grey scale value of the cell free sample (L-DOPA +DMSO) was subtracted from each sample and the MKV3 untreated sample value was set to 1.

**Zebrafish assays**

Maintenance and crossbreeding of WT AB Strain and ctr1 heterozygous Zebrafish: Zebrafish of the WT AB strain and ctr1 heterozygous mutants were bred and maintained following standard protocols. Fertilized embryos were collected from crosses of either WT or *ctr1*$^{+/-}$ x *ctr1*$^{+/-}$ pairs. Embryo treatment and development: For each treatment group, 30 embryos were sorted into individual petri dishes containing 30 mL of embryo buffer supplemented with methylene blue. These embryos were allowed to develop for 3 hours post-fertilization (hpf) before being treated with the specified concentrations of MKV3 or DMSO. The treated embryos were then maintained in an incubator set at 28.5°C until imaging at 48 hpf. Imaging and analysis: Zebrafish larvae were mounted in 3% methylcellulose and imaged using a Zeiss Axioplan 2 microscope equipped with differential interference contrast (DIC) optics at a magnification of 40X. Imaging was performed using an Infinity 5 Teledyne Lumenra camera. Ethical Approval: All experiments involving adult zebrafish were conducted in accordance with the guidelines and regulations approved by the Texas A&M University Institutional Animal Care and Use Committee (IACUC), under protocol no: 2021-0617.

**B16-F10 metastatic melanoma model**

C57BL/6 male mice (age 8 weeks) were inoculated via tail vein with $2.5 \times 10^5$ B16-F10 melanoma cells in sterile PBS. Ten days after inoculation, mice received daily subcutaneous injections of MKV3 (50 mg/kg) or vehicle (5% DMSO / 5% Tween-80) for seven consecutive days. At the end of treatment, mice were euthanized, and lungs were collected, fixed in Bouin's solution, and imaged using a Leica M205 FA stereomicroscope to assess melanin pigmentation of metastatic nodules.

**Growth conditions for etiolated *Arabidopsis* seedlings**

*Arabidopsis thaliana* (ecotype Columbia-0) seeds were surface-sterilized and plated on half-strength Murashige and Skoog (MS) medium as previously described (11), with the following modifications. Filter-sterilized DMSO (vehicle control), 10 µM 1-aminocyclopropane-1-carboxylate (ACC), or the indicated concentration of MKV3 were added to the medium prior to solidification. Seeds were stratified at 4°C for 2 days in darkness and then grown at 22°C for 7 days, wrapped in three layers of aluminum foil to block all light and ensure complete etiolation. Plates were unwrapped after 7 days, and seedlings were immediately imaged and measured.

**Animals**

All mouse procedures conducted in this study were approved by the University of Missouri's Animal Care and Use Committee. Male C57BL/6 mice were obtained from the Jackson Laboratory and were housed in a facility with a 12-hour light-dark cycle with ad libitum access to food and water. The mice were provided with Picolab diet 5053 (containing 13 ppm Cu; PMI International).

**Statistical analysis**

Data are presented as a minimum of three biological replicates and expressed as mean ± standard error of the mean (SEM). Statistical analyses were performed using GraphPad Prism 10.0 software. Data were analyzed using standard Student's *t* test and were considered significant when $p$ values ≤ 0.05. Statistical significance representations: *$p < 0.05$, **$p < 0.01$, ***$p < 0.001$ and ****$p < 0.0001$.

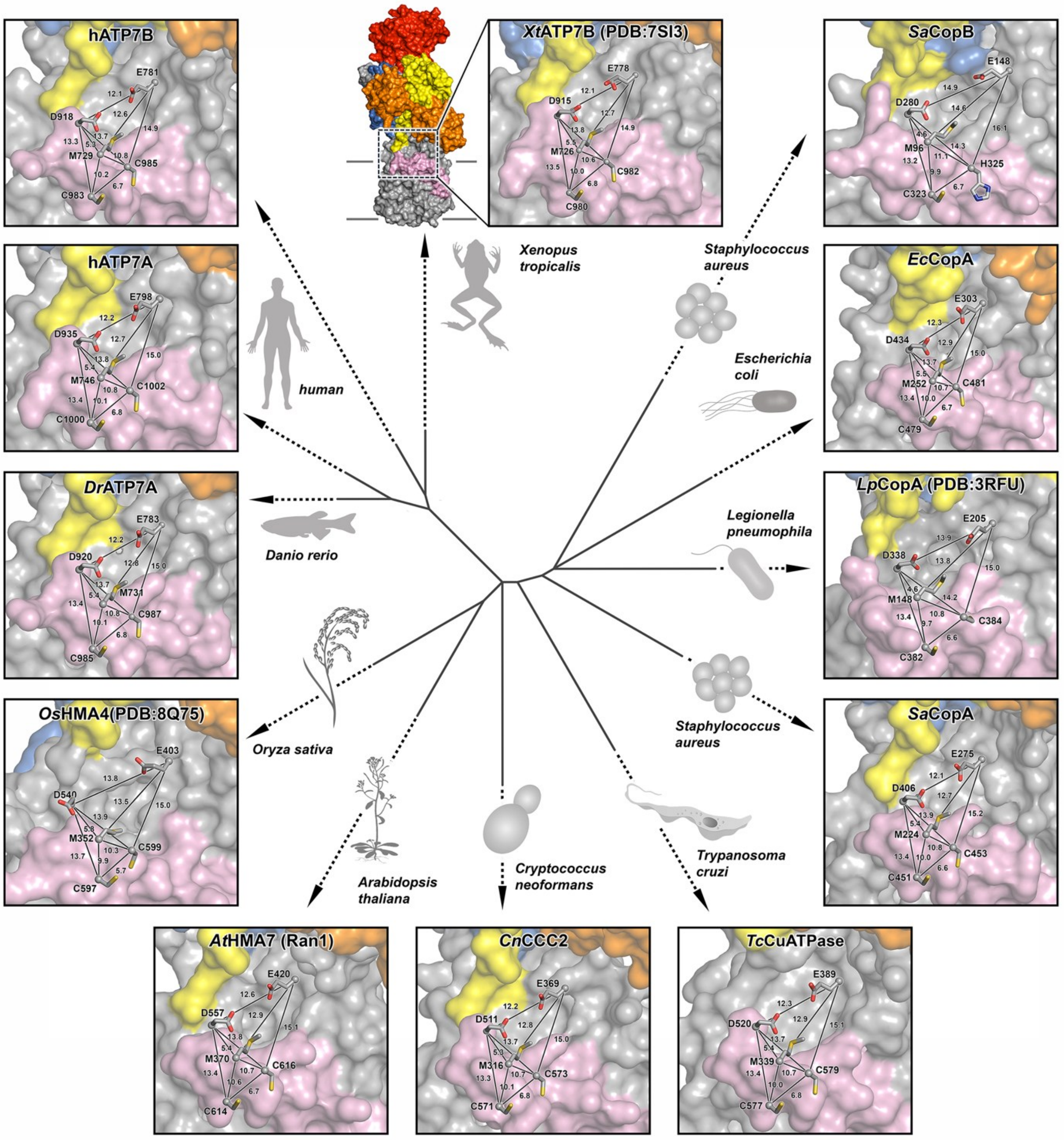


**Figure S1. A conserved druggable pocket at the entrance of $Cu^+$-ATPases.**

$Cu^+$-ATPases in the E2P conformation are shown for *Xenopus tropicalis* ATP7B (PDB 7SI3), *Oryza sativa* HMA4 (PDB 8Q75) and Legionella pneumophila (PDB 3RFU). Structural models were generated from the PDB 7SI3 template using Swiss-Model for human ATP7A and ATP7B, *Danio rerio Dr*ATP7A, *Arabidopsis thaliana At*HMA7, *Cryptococcus neoformans Cn*CCC2, *Trypanosoma cruzi* TcCuATPase, *Staphylococcus aureus Sa*CopA and *Sa*CopB, and *Escherichia coli Ec*CopA. The phylogenetic tree view was generated using Jalview using a Clustal Omega sequence alignment (Fig S2). Distances between resides of the conserved M-E-D triad and cysteines of the CPC motif are shown in Angstroms.

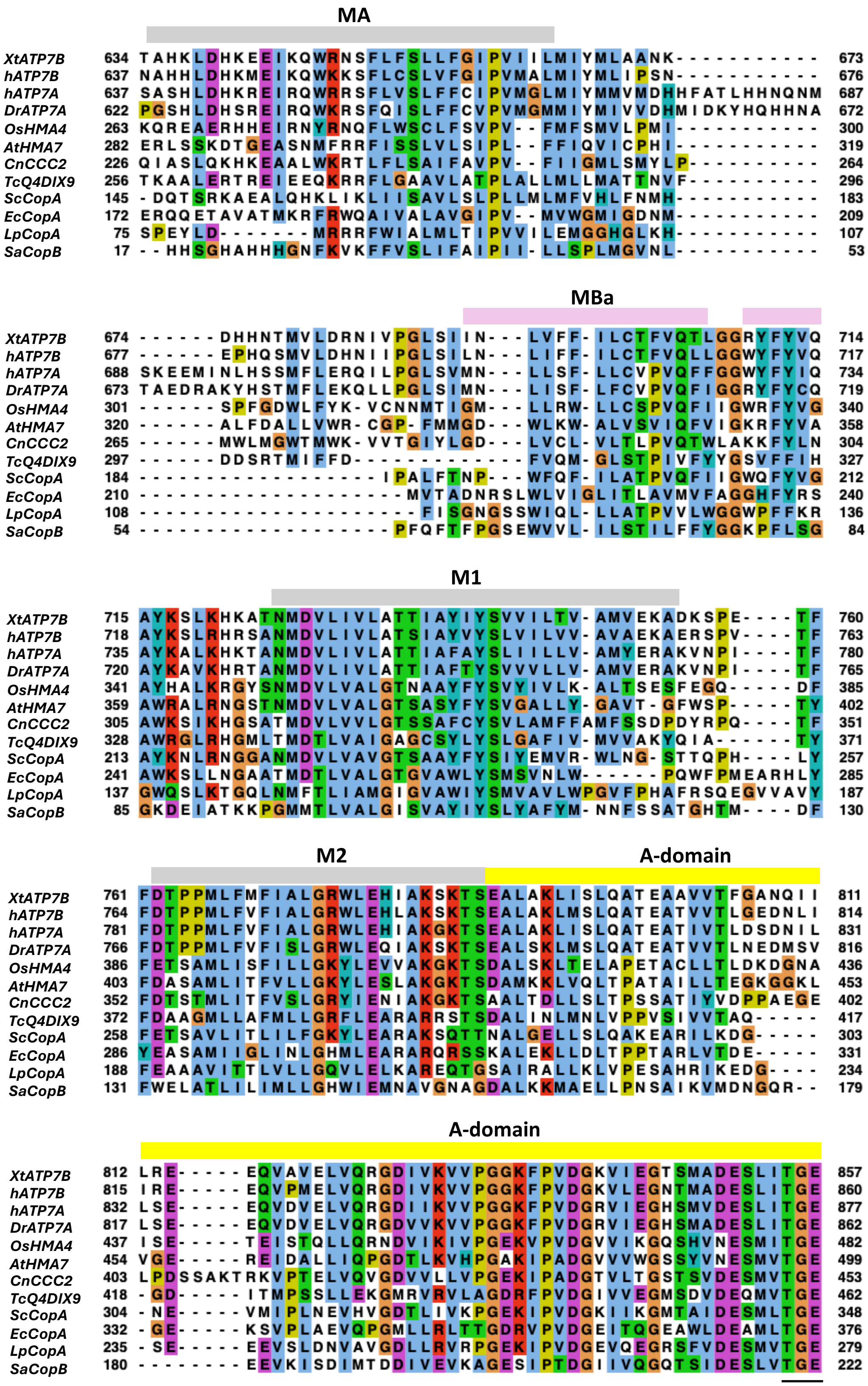
MA
XtATP7B 634 TAHKLDHKEEIKQWRNSFLFSLLFGIPVIILMIYMLAANK----------- 673
hATP7B 637 NAHHLDHKMEIKQWKKSFLCSLVFGIPVMALMIYMLIPSN----------- 676
hATP7A 637 SASHLDHKREIRQWRRSFLVSLFFCIPVMGLMIYMMVMDHHFATLHHNQNM 687
DrATP7A 622 PGSHLDHSREIRQWKRSFQISLFFCVPVMGMMIYMIVVDHMIDKYHQHHNA 672
OsHMA4 263 KQREAERHHEIRNYRNQFLWSCLFSVPV--FMFSMVLPMI----------- 300
AtHMA7 282 ERLSSKDTGEASNMFRRFISSLVLSIPL--FFIQVICPHI----------- 319
CnCCC2 226 QIASLQKHKEAALWKRTLFLSAIFAVPV--FIIGMLSMYLP---------- 264
TcQ4DIX9 256 TKAALERTREIEEQKRRFLGAAVLATPLALLMLLMATTNVF---------- 296
ScCopA 145 -DQTSRKAEALQHKLIKLIISAVLSLPLLMLMFVHLFNMH----------- 183
EcCopA 172 ERQQETAVATMKRFRWQAIVALAVGIPV--MVWGMIGDNM----------- 209
LpCopA 75 SPEYLD--------MRRRFWIALMLTIPVVILEMGGHGLKH---------- 107
SaCopB 17 --HHSGHAHHHGNFKVKFFVSLIFAIPII-LLSPLMGVNL----------- 53
MBa
XtATP7B 674 ------DHHNTMVLDRNIVPGLSIIN---LVFF-ILCTFVQTLGGRYFYVQ 714
hATP7B 677 ------EPHQSMVLDHNIIPGLSILN---LIFF-ILCTFVQLLGGWYFYVQ 717
hATP7A 688 SKEEMINLHSSMFLERQILPGLSVMN---LLSF-LLCVPVQFFGGWYFYIQ 734
DrATP7A 673 TAEDRAKYHSTMFLEKQLLPGLSIMN---LISF-LFCVPVQFIGGRYFYCQ 719
OsHMA4 301 ------SPFGDWLFYK-VCNNMTIGM---LLRW-LLCSPVQFIIGWRFYVG 340
AtHMA7 320 ------ALFDALLVWR-CGP-FMMGD---WLKW-ALVSVIQFVIGKRFYVA 358
CnCCC2 265 ------MWLMGWTMWK-VVTGIYLGD---LVCL-VLTLPVQTWLAKKFYLN 304
TcQ4DIX9 297 ------DDSRTMIFFD-------------FVQM-GLSTPIVFYYGSVFFIH 327
ScCopA 184 ------------------IPALFTNP---WFQF-ILATPVQFIIGWQFYVG 212
EcCopA 210 --------------------MVTADNRSLWLVIGLITLAVMVFAGGHFYRS 240
LpCopA 108 ---------------------FISGNGSSWIQL-LLATPVVLWGGWPFFKR 136
SaCopB 54 -------------------PFQFTFPGSEWVVL-ILSTILFFYGGKPFLSG 84
M1
XtATP7B 715 AYKSLKHKATNMDVLIVLATTIAYIYSVVILTV-AMVEKADKSPE----TF 760
hATP7B 718 AYKSLRHRSANMDVLIVLATSIAYVYSLVILVV-AVAEKAERSPV----TF 763
hATP7A 735 AYKALKHKTANMDVLIVLATTIAFAYSLIILLV-AMYERAKVNPI----TF 780
DrATP7A 720 AYKAVKHRTANMDVLIVLATTIAFTYSVVVLLV-AMVERAKVNPI----TF 765
OsHMA4 341 AYHALKRGYSNMDVLVALGTNAAYFYSVYIVLK-ALTSESFEGQ-----DF 385
AtHMA7 359 AWRALRNGSTNMDVLVALGTSASYFYSVGALLY-GAVT-GFWSP-----TY 402
CnCCC2 305 AWKSIKHGSATMDVLVVLGTSSAFCYSVLAMFFAMFSSDPDYRPQ----TF 351
TcQ4DIX9 328 AWRGLRHGMLTMDTLVAIGAGCSYLYSLGAFIV-MVVAKYQIA------TY 371
ScCopA 213 AYKNLRNGGANMDVLVAVGTSAAYFYSIYEMVR-WLNG-STTQPH----LY 257
EcCopA 241 AWKSLLNGAATMDTLVALGTGVAWLYSMSVNLW------PQWFPMEARHLY 285
LpCopA 137 GWQSLKTGQLNMFTLIAMGIGVAWIYSMVAVLWPGVFPHAFRSQEGVVAVY 187
SaCopB 85 GKDEIATKKPGMMTLVALGISVAYIYSLYAFYM-NNFSSATGHTM----DF 130
M2
A-domain
XtATP7B 761 FDTPPMLFMFIALGRWLEHIAKSKTSEALAKLISLQATEAAVVTFGANQII 811
hATP7B 764 FDTPPMLFVFIALGRWLEHLAKSKTSEALAKLMSLQATEATVVTLGEDNLI 814
hATP7A 781 FDTPPMLFVFIALGRWLEHIAKGKTSEALAKLISLQATEATIVTLDSDNIL 831
DrATP7A 766 FDTPPMLFVFISLGRWLEQIAKSKTSEALSKLMSLQATEATVVTLNEDMSV 816
OsHMA4 386 FETSAMLISFILLGKYLEVVAKGKTSDALSKLTELAPETACLLTLDKDGNA 436
AtHMA7 403 FDASAMLITFVLLGKYLESLAKGKTSDAMKKLVQLTPATAILLTEGKGGKL 453
CnCCC2 352 FDTSTMLITFVSLGRYIENIAKGKTSAALTDLLSLTPSSATIYVDPPAEGE 402
TcQ4DIX9 372 FDAAGMLLAFMLLGRFLEARARRSTSDALINLMNLVPPVSIVVTAQ----- 417
ScCopA 258 FETSAVLITLILFGKYLEARAKSQTTNALGELLSLQAKEARILKDG----- 303
EcCopA 286 YEASAMIIGLINLGHMLEARARQRSSKALEKLLDLTPPTARLVTDE----- 331
LpCopA 188 FEAAAVITTLVLLGQVLELKAREQTGSAIRALLKLVPESAHRIKEDG---- 234
SaCopB 131 FWELATLILIMLLGHWIEMNAVGNAGDALKKMAELLPNSAIKVMDNGQR-- 179
A-domain
XtATP7B 812 LRE-----EQVAVELVQRGDIVKVVPGGKFPVDGKVIEGTSMADESLITGE 857
hATP7B 815 IRE-----EQVPMELVQRGDIVKVVPGGKFPVDGKVLEGNTMADESLITGE 860
hATP7A 832 LSE-----EQVDVELVQRGDIIKVVPGGKFPVDGRVIEGHSMVDESLITGE 877
DrATP7A 817 LSE-----EQVDVELVQRGDVVKVVPGGKFPVDGRVIEGHSMADESLITGE 862
OsHMA4 437 ISE-----TEISTQLLQRNDVIKIVPGEKVPVDGVVIKGQSHVNESMITGE 482
AtHMA7 454 VGE-----REIDALLIQPGDTLKVHPGAKIPADGVVVWGSSYVNESMVTGE 499
CnCCC2 403 LPDSSAKTRKVPTELVQVGDVVLLVPGEKIPADGTVLTGSTSVDESMVTGE 453
TcQ4DIX9 418 -GD-----ITMPSSLLEKGMRVRVLAGDRFPVDGIVVEGMSDVDEQMVTGE 462
ScCopA 304 -NE-----VMIPLNEVHVGDTLIVKPGEKIPVDGKIIKGMTAIDESMLTGE 348
EcCopA 332 -GE-----KSVPLAEVQPGMLLRLTTGDRVPVDGEITQGEAWLDEAMLTGE 376
LpCopA 235 -SE-----EEVSLDNVAVGDLLRVRPGEKIPVDGEVQEGRSFVDESMVTGE 279
SaCopB 180 --------EEVKISDIMTDDIVEVKAGESIPTDGIIVQGQTSIDESLVTGE 222

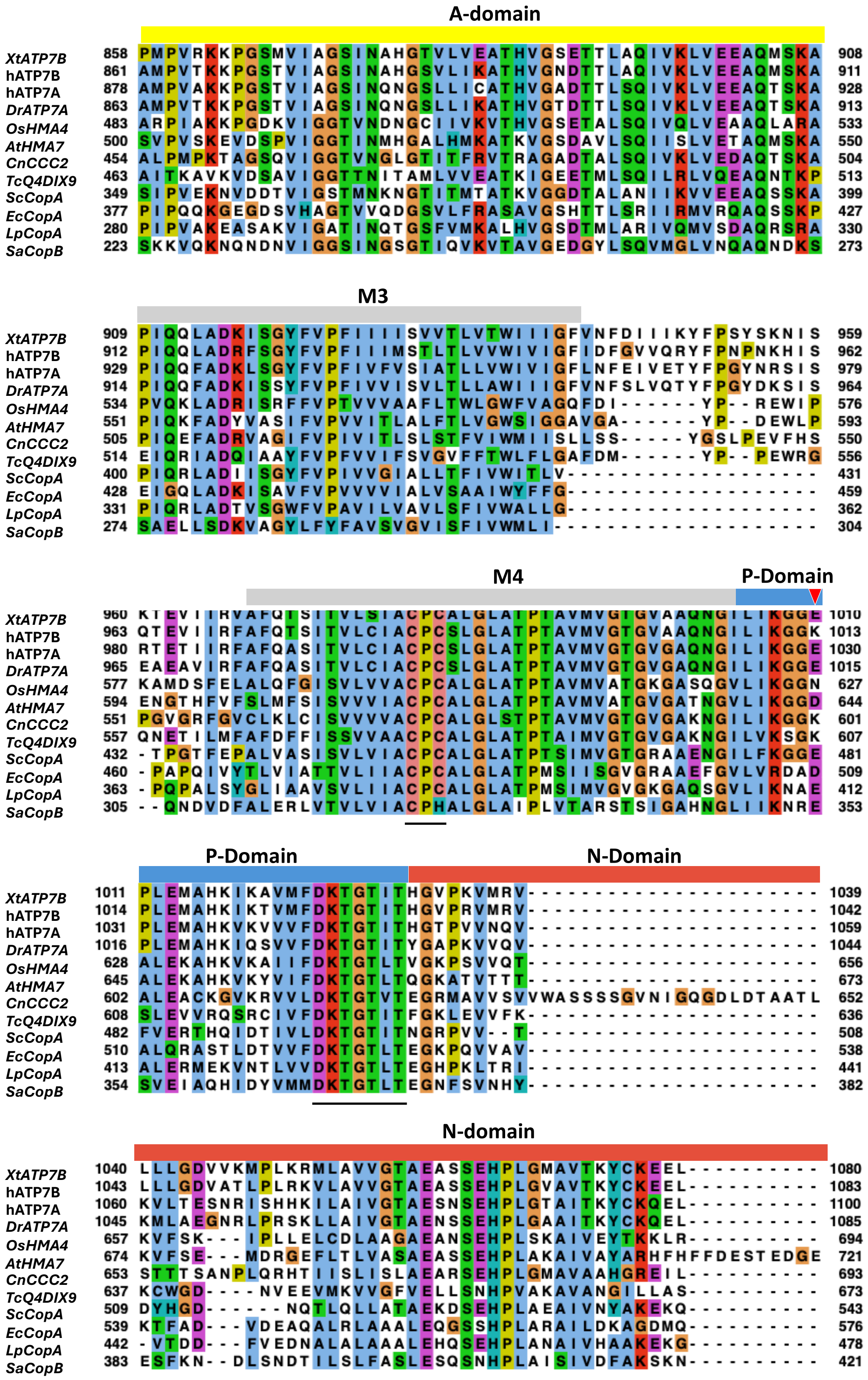
A-domain
M3
M4
P-Domain
P-Domain
N-Domain
N-domain
XtATP7B
hATP7B
hATP7A
DrATP7A
OsHMA4
AtHMA7
CnCCC2
TcQ4DIX9
ScCopA
EcCopA
LpCopA
SaCopB

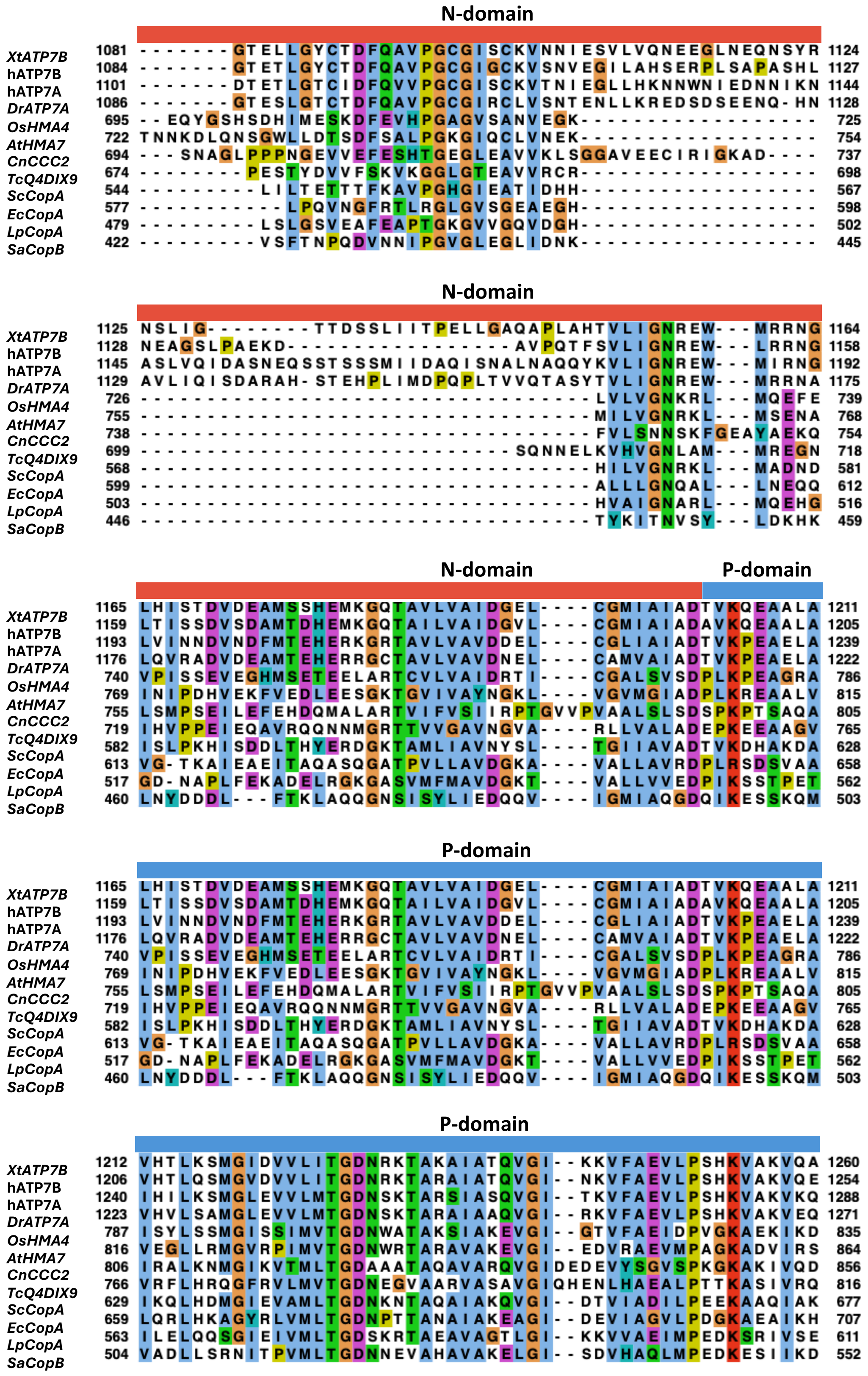
N-domain
N-domain
N-domain
P-domain
P-domain
P-domain
XtATP7B
hATP7B
hATP7A
DrATP7A
OsHMA4
AtHMA7
CnCCC2
TcQ4DIX9
ScCopA
EcCopA
LpCopA
SaCopB

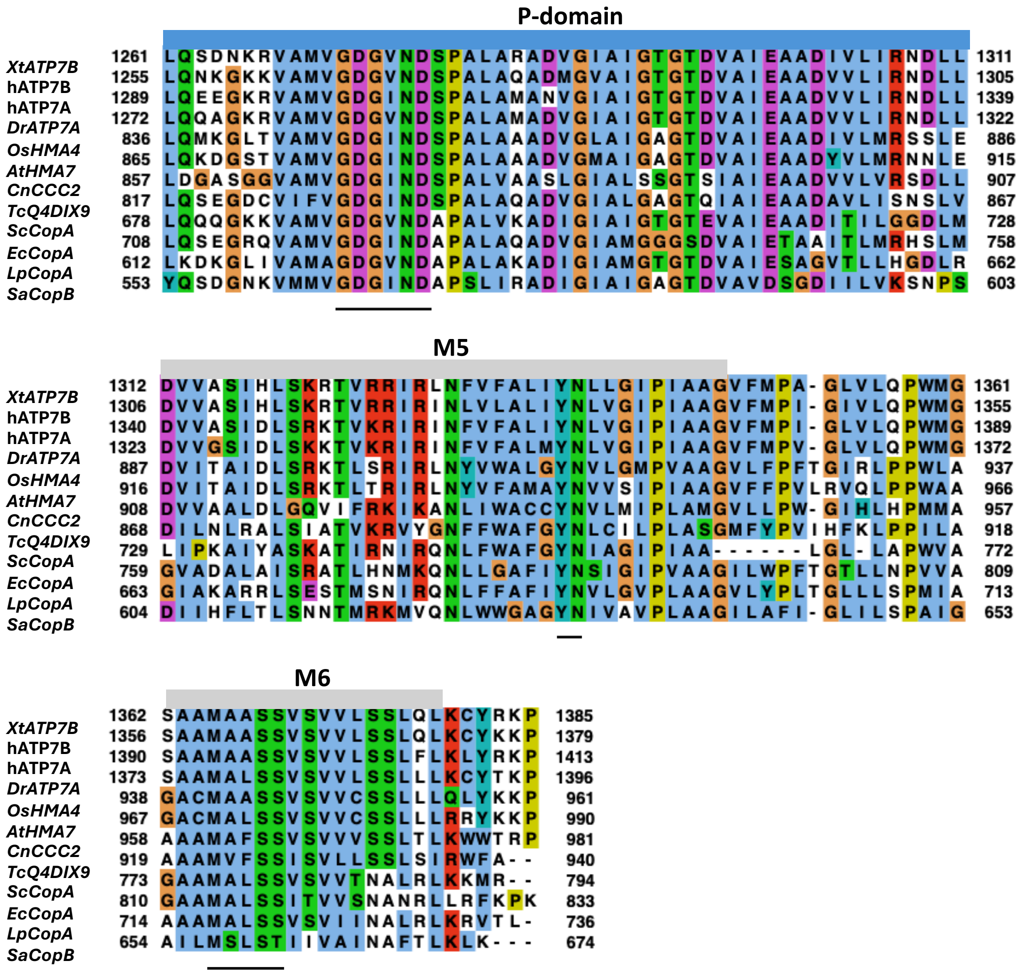


**Figure S2. Conserved features among copper transporting P-type ATPases.** The alignment spans the first (MA) to the eighth (M6) transmembrane domains for the following Cu-ATPases: *Xenopus tropicalis* ATP7B (A0A6I8R0A5), human ATP7B (P35670), human ATP7A (Q04656), *Danio rerio DrATP7A* (Q4F8H5), *Oryza sativa Os*HMA4 (Q6H7M3), *Arabidopsis thaliana* HMA7 (RAN1; Q9S7J8), *Cryptococcus neoformans Cn*CCC2 (Q5K722), *Trypanosoma cruzi TcCuATPase* (Q4DIX9), *Staphylococcus aureus Sa*CopA (Q2FV64), *Escherichia coli Ec*CopA (Q59385), *Legionella pneumophila Lp*CopA (Q5ZWR1), *Staphylococcus aureus Sa*CopB (Q2FKI2). Uniprot numbers are shown in parentheses. Alignments were generated using Clustal Omega and visualized using Jalview. Conserved signature sequences are underlined including the phosphatase motif (TGE) in the A-domain, the aspartyl-phosphate (DKTGTI/LT) and ATP binding motifs GDG(I/V)ND in the P-domain, the copper-binding CPC/H motif in M4, YN in M5, and MXXSS in M6.

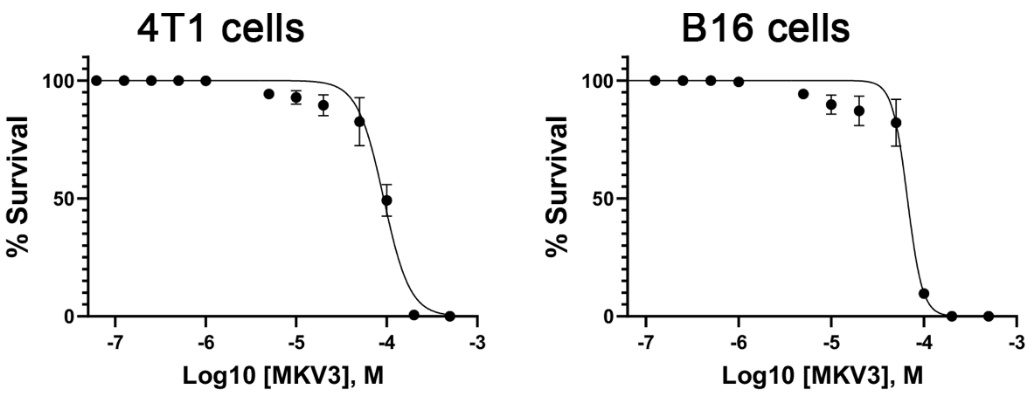


**Figure S3. Survival of 4T1 and B16 cells in various concentrations of MKV3.** Survival was measured after 48 h using the Crystal Violet assay. The $CC_{50}$ value (mean ± SEM) for 4T1 cells was 92.5 ± 14 μM, and for B16 cells was 66.2 ± 6 μM.

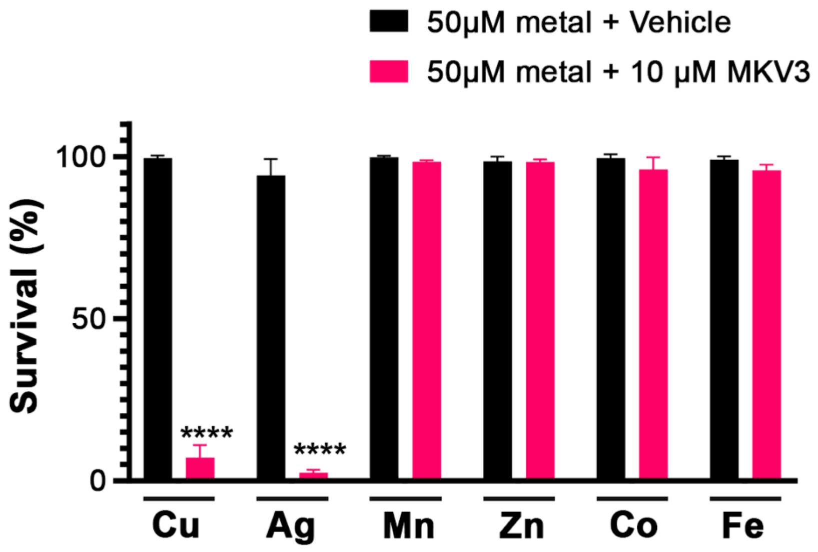


**Figure S4. MKV3 potentiates sensitivity to copper and silver, but not other heavy metals.** Survival of 4T1 cells was measured using the Crystal Violet assay after a 48-hour treatment with 10 μM MKV3 or 1% DMSO (vehicle) alone or in combination with 50 μM of the indicated metals (added as $CuCl_2$, $AgNO_3$, $MnCl_2$, $ZnCl_2$, $CoCl_2$ or $FeCl_2$). Data are expressed as percent survival against the vehicle control for each metal (mean ± SEM; ****$p < 0.0001$).

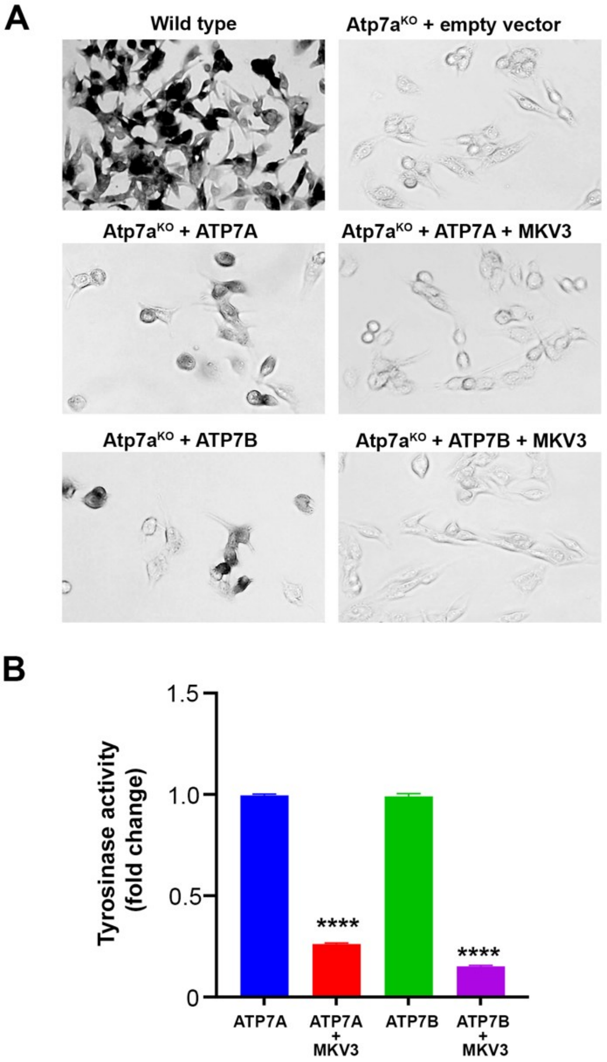


**Figure S5: MKV3 inhibits ATP7A- and ATP7B-dependent restoration of tyrosinase in *Atp7a*-knockout B16-F10 melanoma cells.** (A) *In situ* tyrosinase activity was evaluated in wild-type B16 melanoma cells and *Atp7a*-knockout (Atp7a$^{KO}$) cells transiently transfected with human ATP7A or ATP7B, followed by treatment with 10 μM MKV3 for 24 hours or left untreated. Representative images of pigmented cells under each condition are shown. (B) Quantitative analysis of tyrosinase activity with or without MKV3 treatment (mean ±SEM; ****$p < 0.0001$).

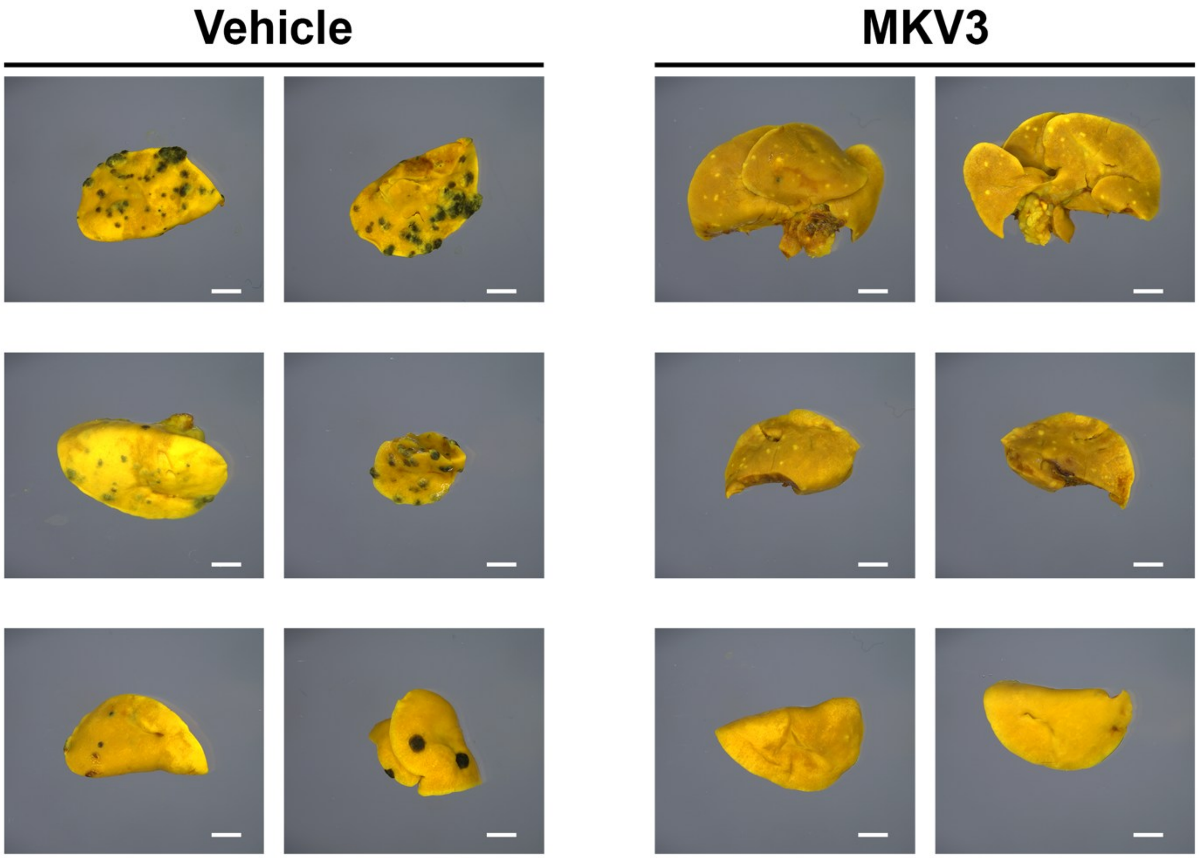


**Figure S6. MKV3 reduces melanogenesis within metastatic melanoma tumors in mice.** Images of lung lobes derived from B16-F10 tumor-bearing mice treated subcutaneously with either vehicle control (5% v/v DMSO /Tween-80) or MKV3 (50 mg/kg) for 7 days. Lungs were fixed, stained with Bouin's solution, and imaged using a Leica M205 FA stereomicroscope. Scale bars = 2mm.

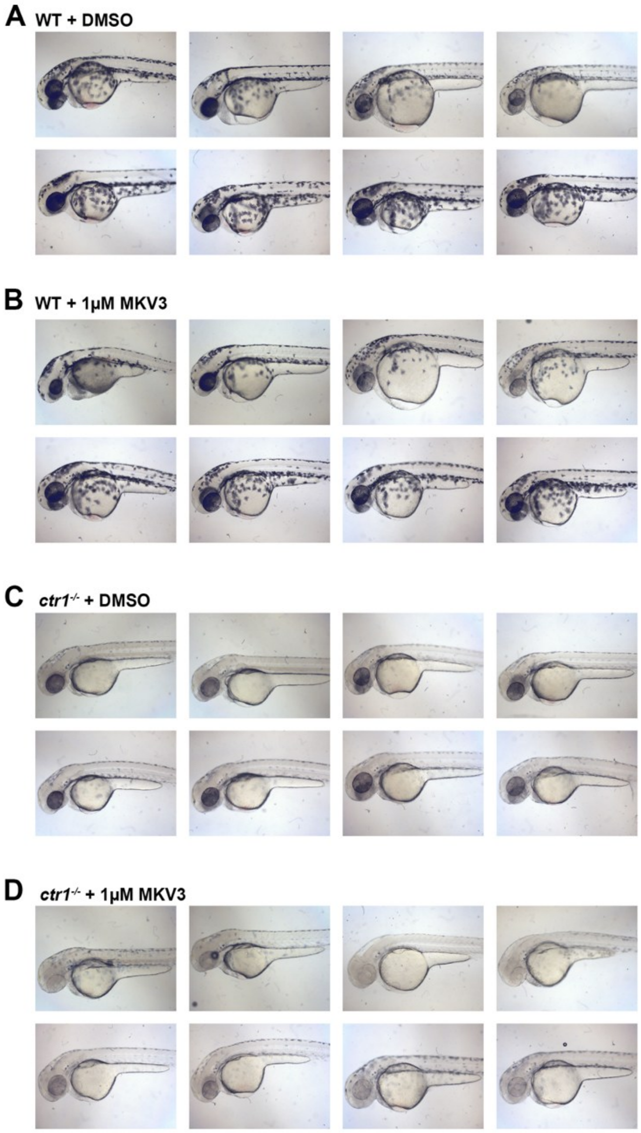


**Figure S7. MKV3 reduces melanogenesis in *ctr1*$^{-/-}$ zebrafish embryos.** Bright-field images of WT and *ctr1*$^{-/-}$ zebrafish embryos after 48 hours treatment with either MKV3 (1 μM) or DMSO (vehicle control).

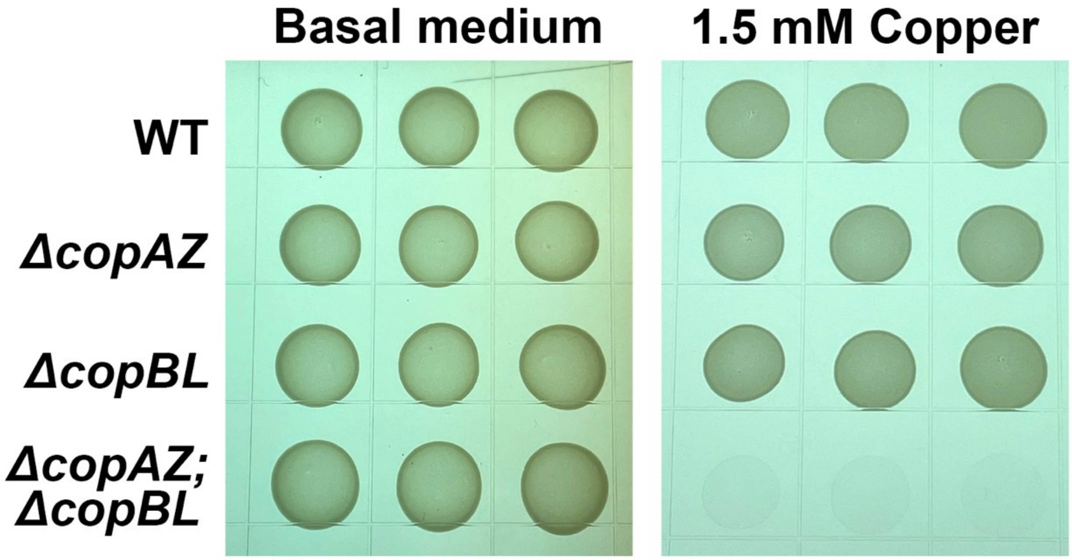


**Figure S8. Growth of USA300 MRSA variants on NB1 agar plates with and without supplemental copper.** Equivalent amounts of MRSA strains were spotted in triplicate onto NB1 agar plates with or without 1.5 mM $CuSO_4$ and incubated overnight at 37$^{o}$C.

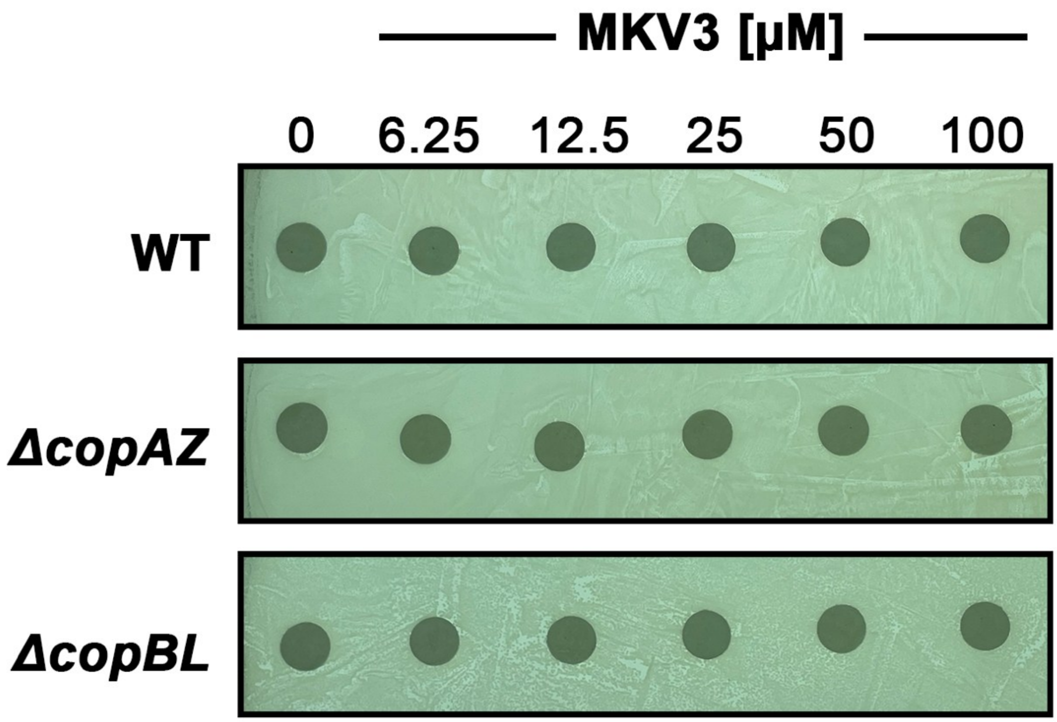


**Figure S9. MKV3 zone of inhibition assay on NB1 agar without added copper.** Filter paper discs containing the indicated concentrations of MKV3 were placed onto WT, *ΔcopAZ*, and Δ*copBL* strains of MRSA on NB1 agar medium without added copper and incubated overnight at 37$^{o}$C.

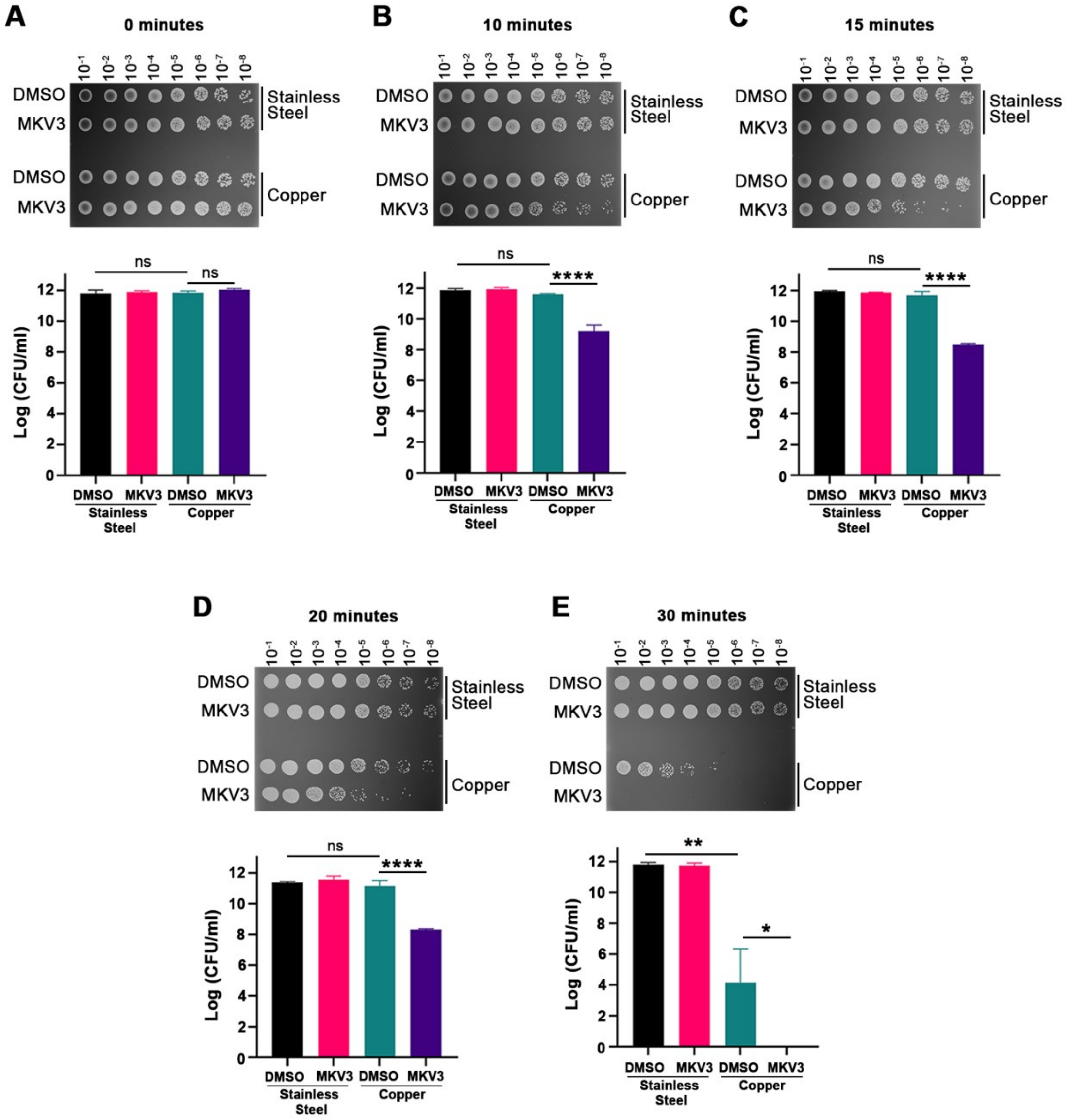


**Figure S10. Contact killing of MRSA on solid copper is potentiated by MKV3 in a time-dependent manner.** MRSA was applied to stainless-steel or copper surfaces and exposed for 0–30 minutes in the presence of 10 µM MKV3 or DMSO control. (A–E) Bacterial spot-assay images and CFU quantification are shown for each time point. Data represent mean ± SEM, with statistical analyses indicated.

A

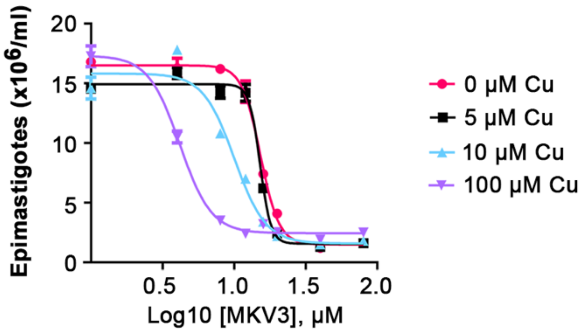


B

| Copper (μM) | MKV3 $CC_{50}$ (μM) |
|---|---|
| 0 | 15.25 ± 0.26 |
| 5 | 14.69 ± 1.24 |
| 10 | 10.63 ± 0.80*** |
| 100 | 4.47 ± 0.28**** |

C

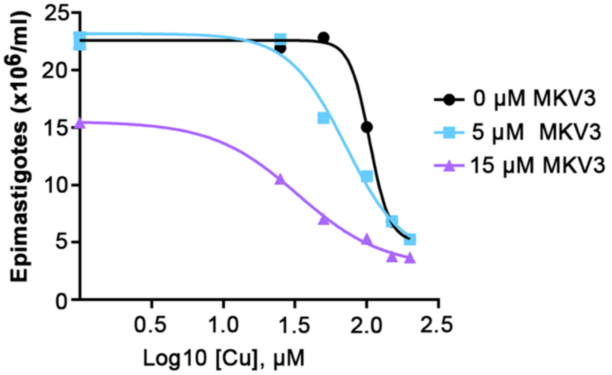


D

| MKV3 (μM) | Cu $CC_{50}$ (μM) |
|---|---|
| 0 | 108.17 ± 1.81 |
| 5 | 70.45 ± 2.63 ***** |
| 15 | 32.97 ± 0.42 ***** |

**Figure S11. Dose-response curves of MKV3 and copper in *T. cruzi* epimastigotes.** (A) *T. cruzi* epimastigotes were quantified as a function of MKV3 dose at the indicated copper concentrations. (B) The $CC_{50}$ values for MKV3 at each copper concentration in panel A. (C) *T. cruzi* epimastigotes were quantified as a function of copper concentration at the indicated doses of MKV3. (D) The $CC_{50}$ values for copper at each MKV3 dose in panel C.

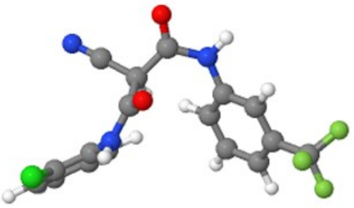

MKV2

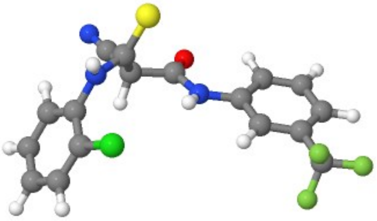

MKV3

**Figure S12. Geometry-optimized conformers of MKV2 and MKV3 calculated using *ab initio* quantum chemical methods (6-31G* basis set, $P_{SI}$4) (12).** Optimization reveals distinct low-energy conformations for MKV2 and MKV3.

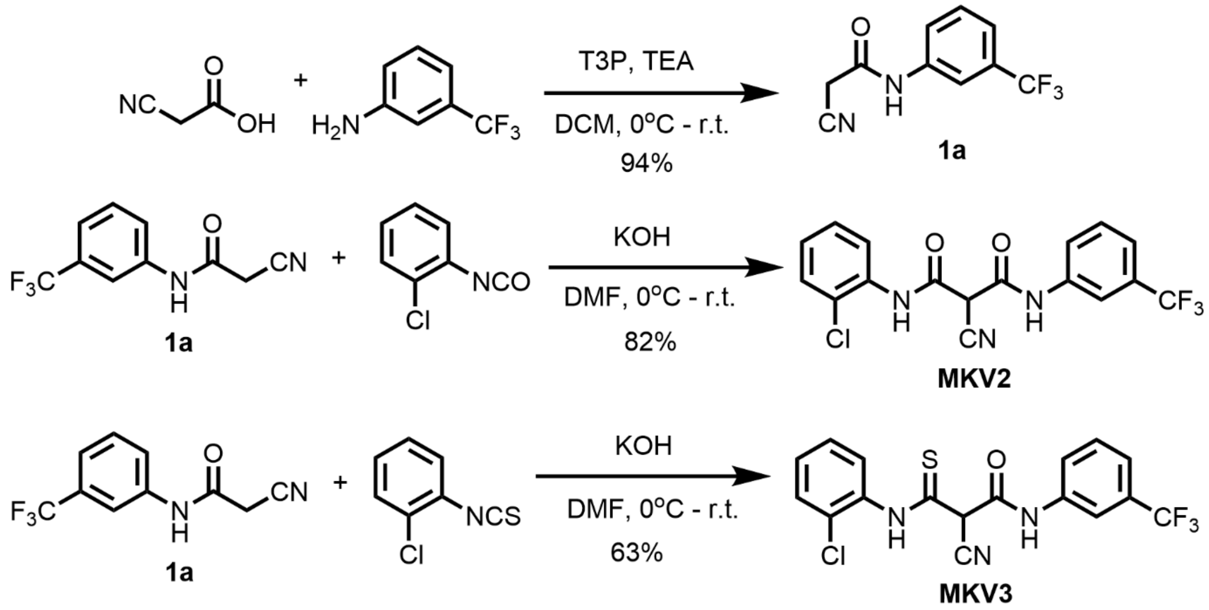


**Figure S13.** Schemes for the synthesis of **MKV2** and **MKV3.**

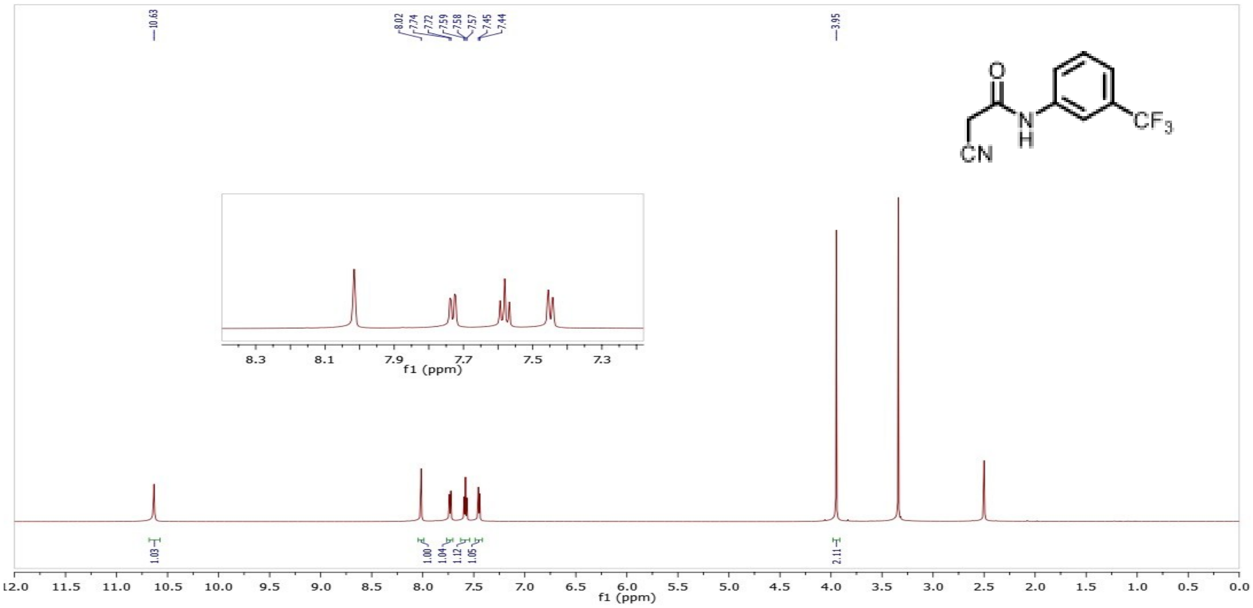


**Figure S14**. $^{1}$H NMR spectra of Intermediate **1a** in DMSO

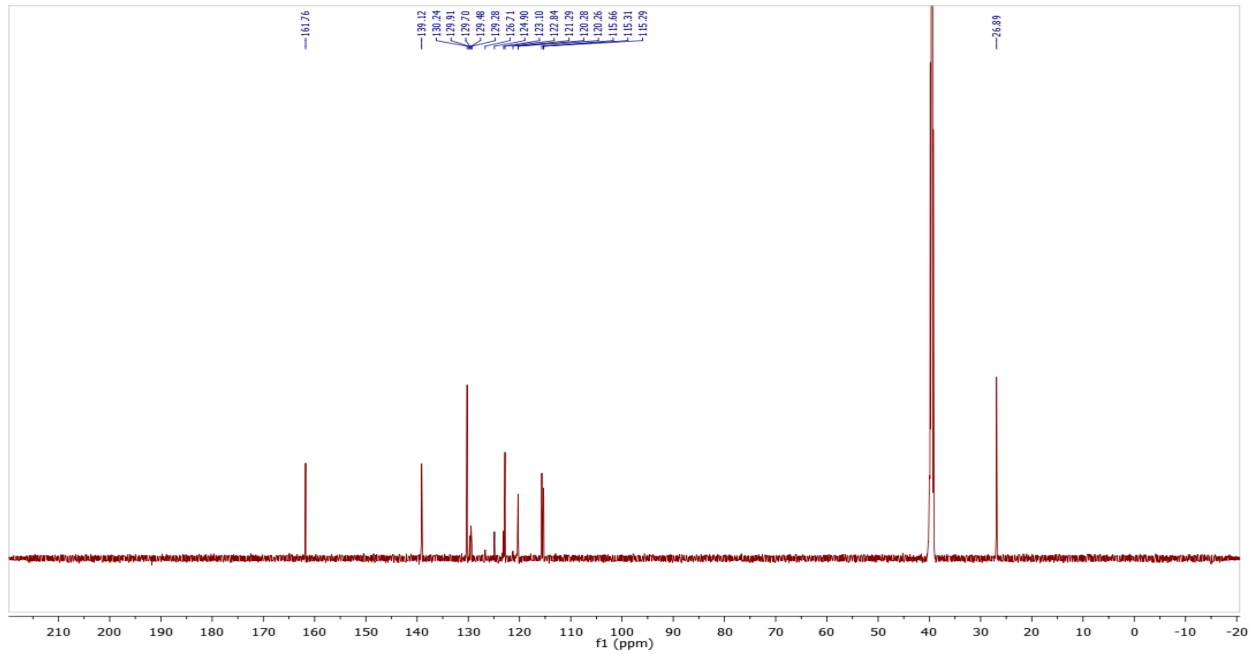


**Figure S15**. $^{13}C$ NMR spectra of Intermediate **1a** in DMSO

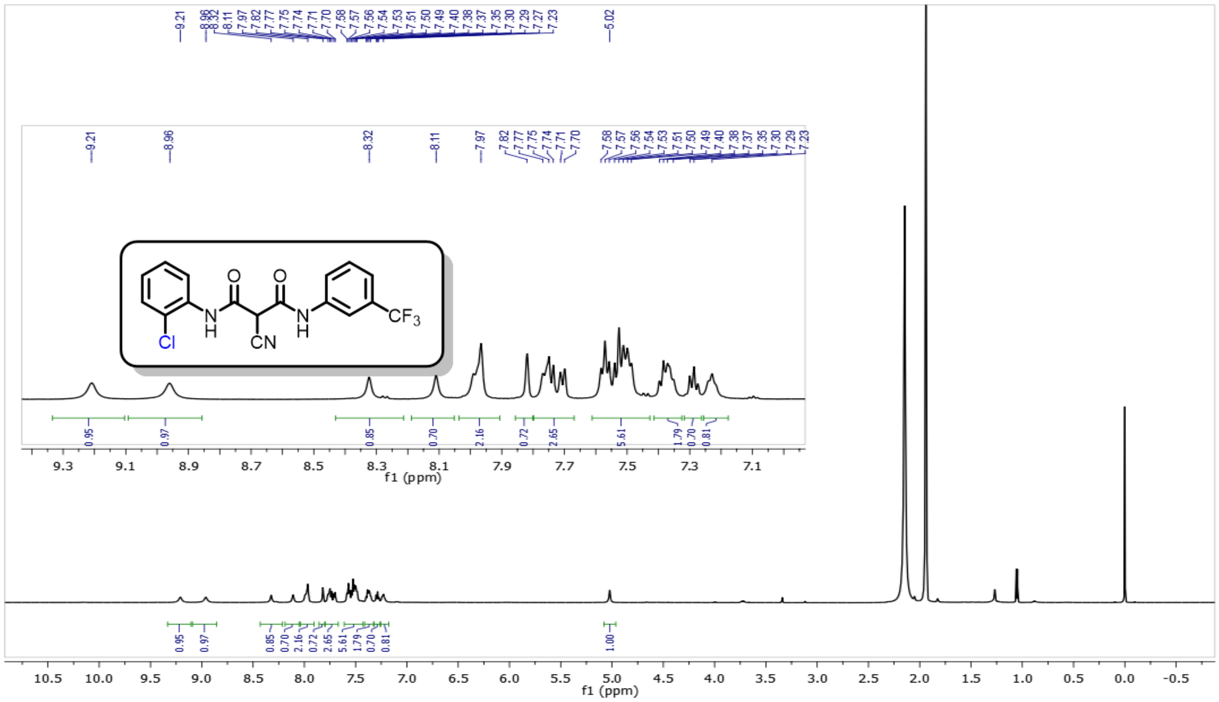

**Figure S16**.[1]H NMR of **MKV2** in $CD_3CN$ (**MKV2** was obtained as a mixture of keto and enol tautomers)

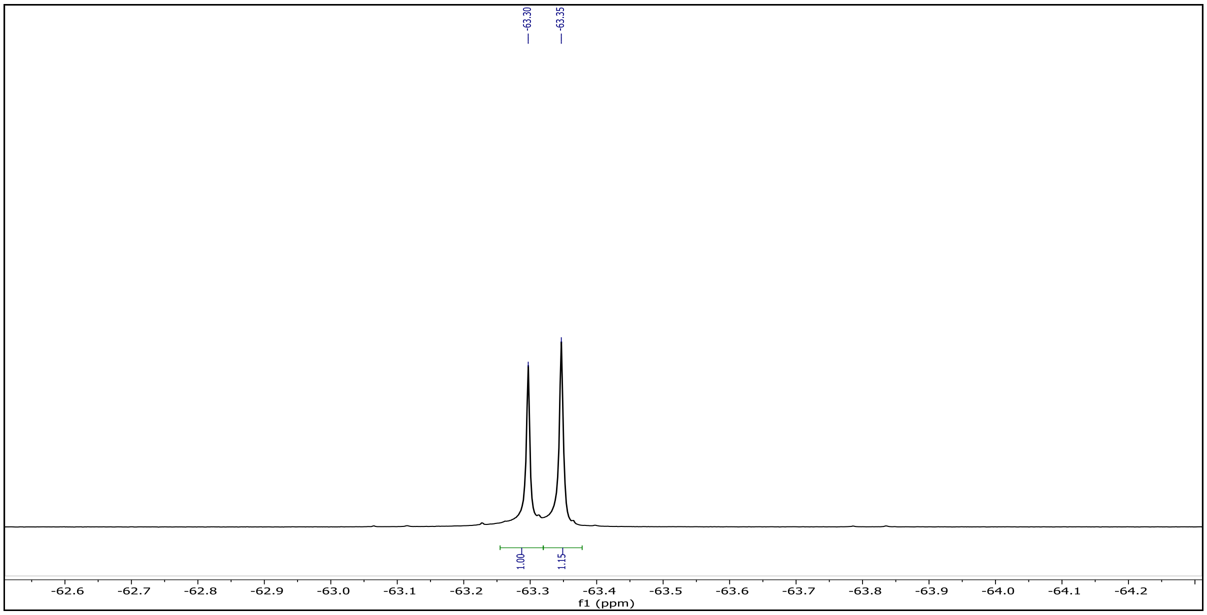


**Figure S17**. $^{19}F$ NMR of **MKV2** in $CD_3CN$ shows the presence of signals from keto and enol tautomers

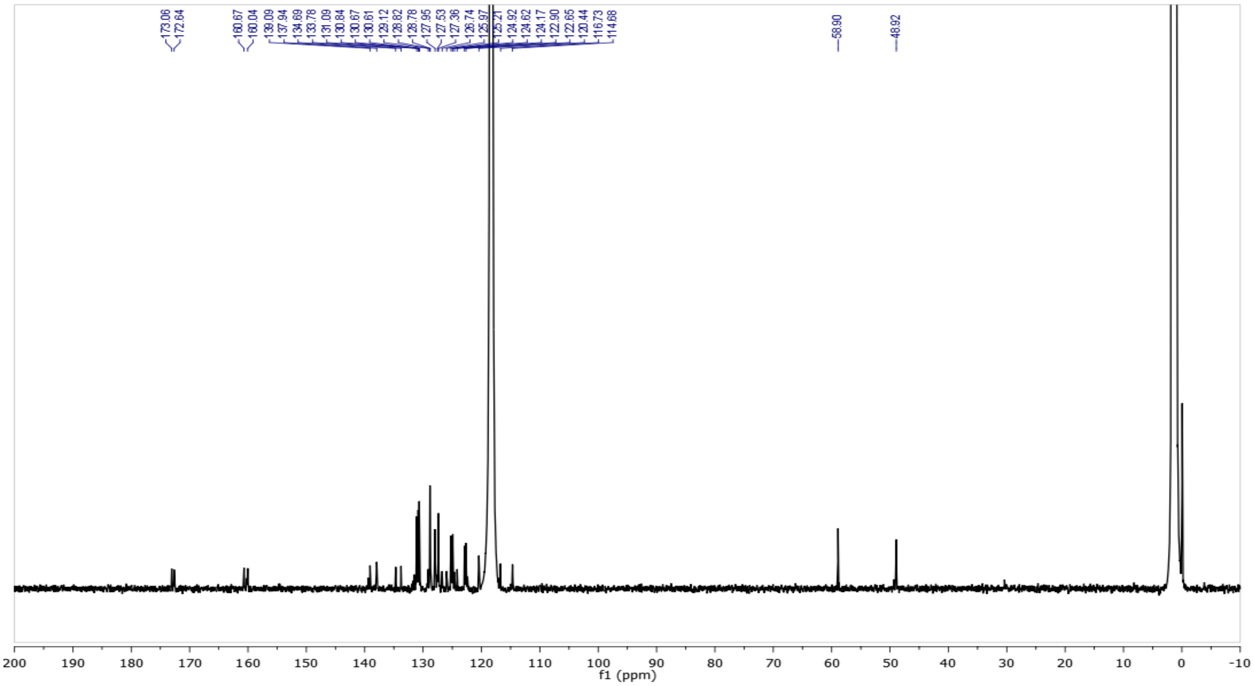


**Figure S18.** $^{13}C$ NMR of **MKV2** in $CD_3CN$

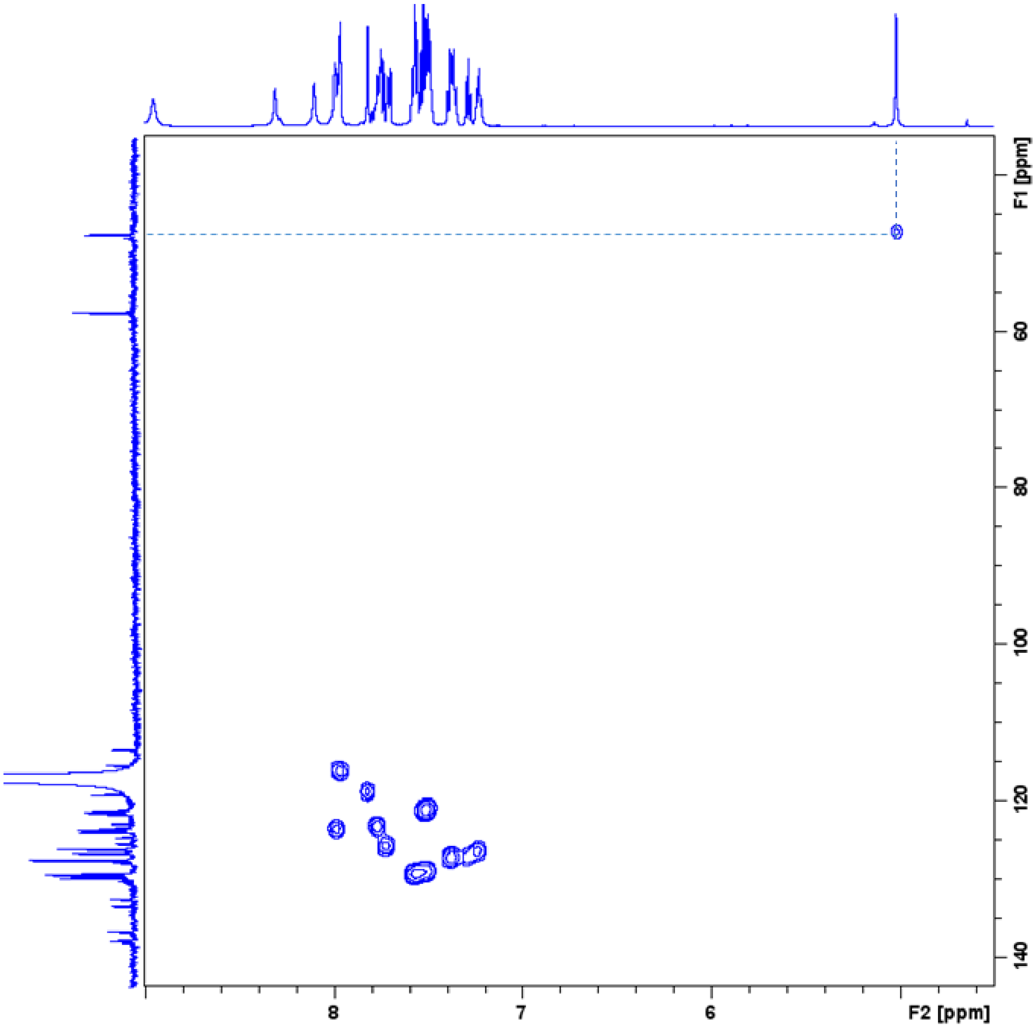


**Figure S19**. HSQC spectrum of **MKV2** in $CD_3CN$. **MKV2** was obtained as a mixture of keto and enol tautomers. The 1-bond correlation mapped out here shows the interaction of the methine group attached to the nitrile group in the keto form to its respective carbon signal.

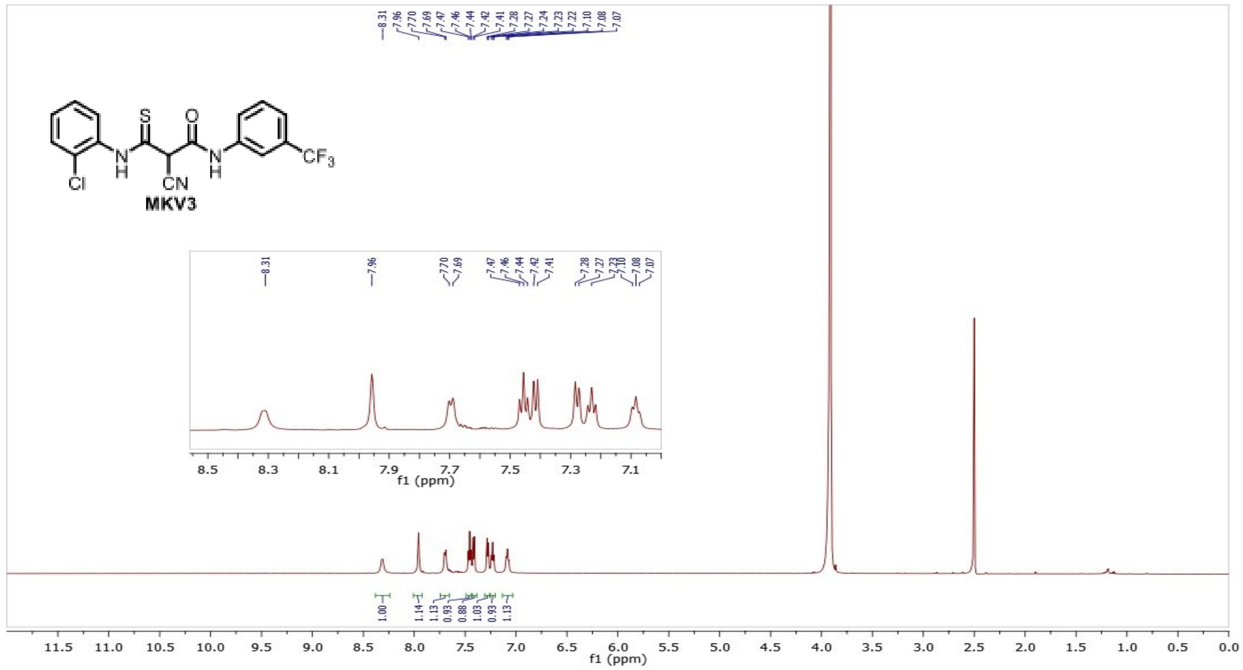


**Figure S20**. $^{1}H$ NMR spectra of **MKV3** in (10: 1) DMSO-$d_6$: $D_2O$

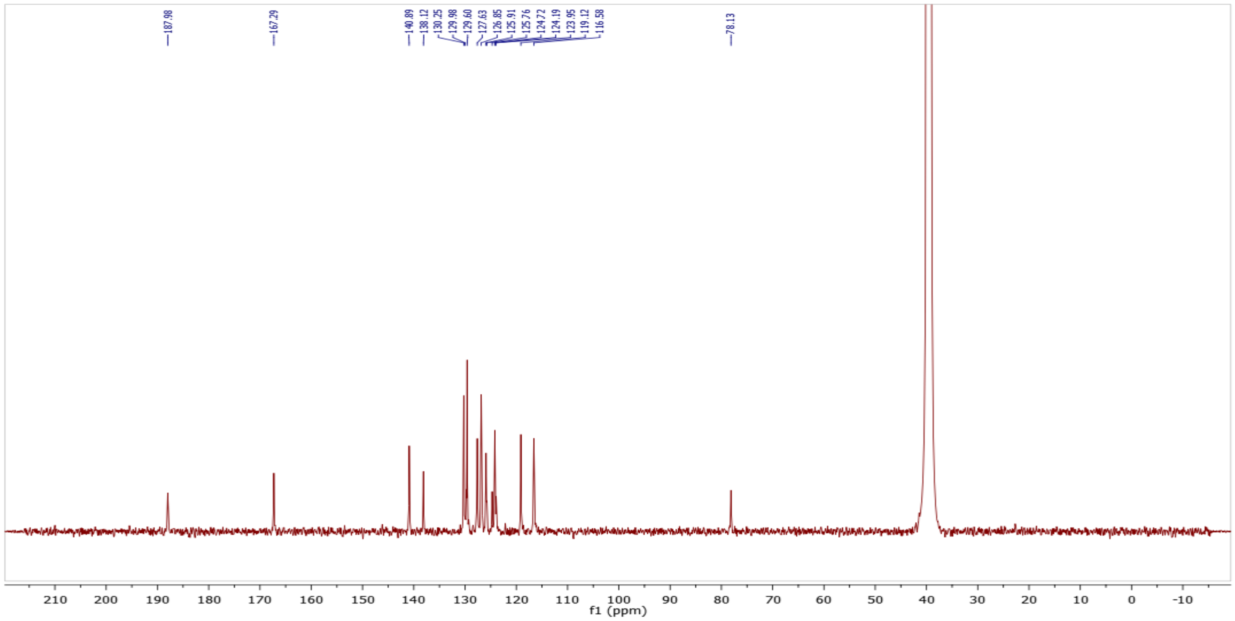


**Figure S21**. [13]C NMR spectra of **MKV3** in (10: 1) DMSO-$d_6$: $D_2O$

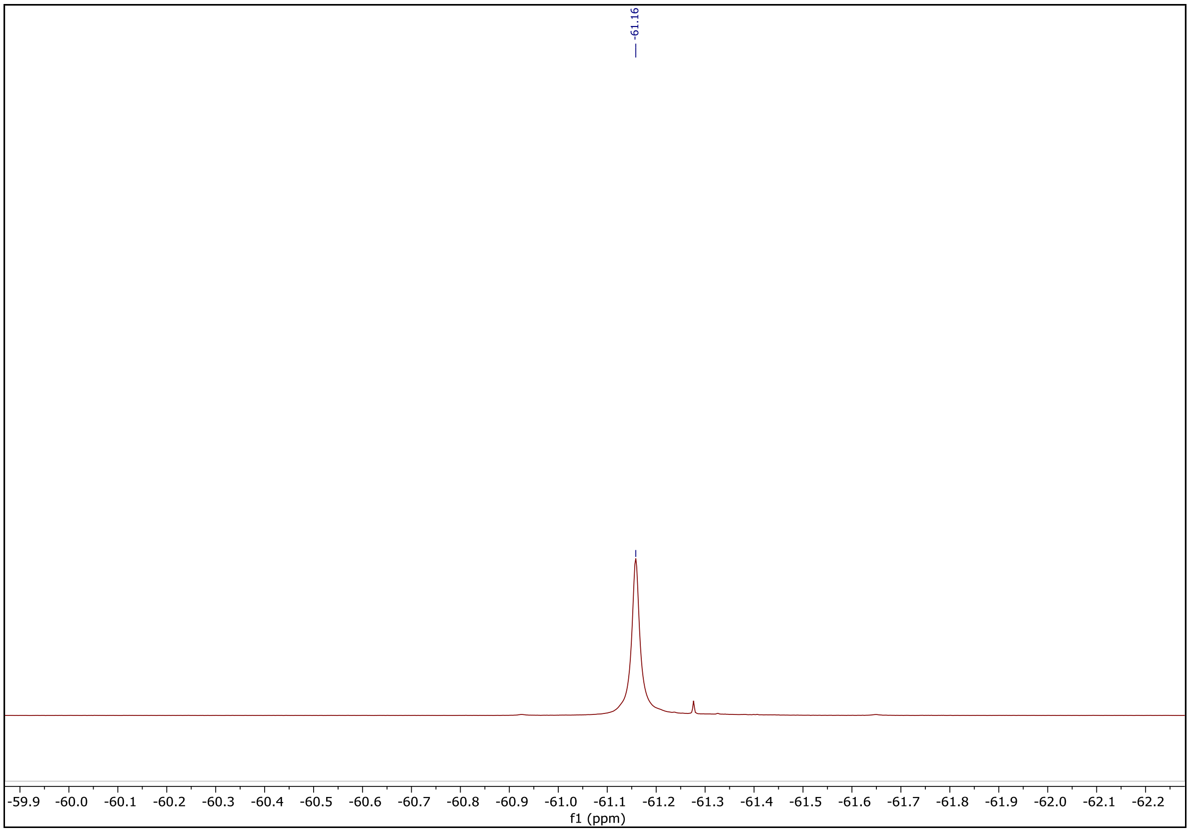


**Figure S22**. $^{19}F$ NMR spectra of **MKV3** in (10: 1) DMSO-$d_6$: $D_2O$

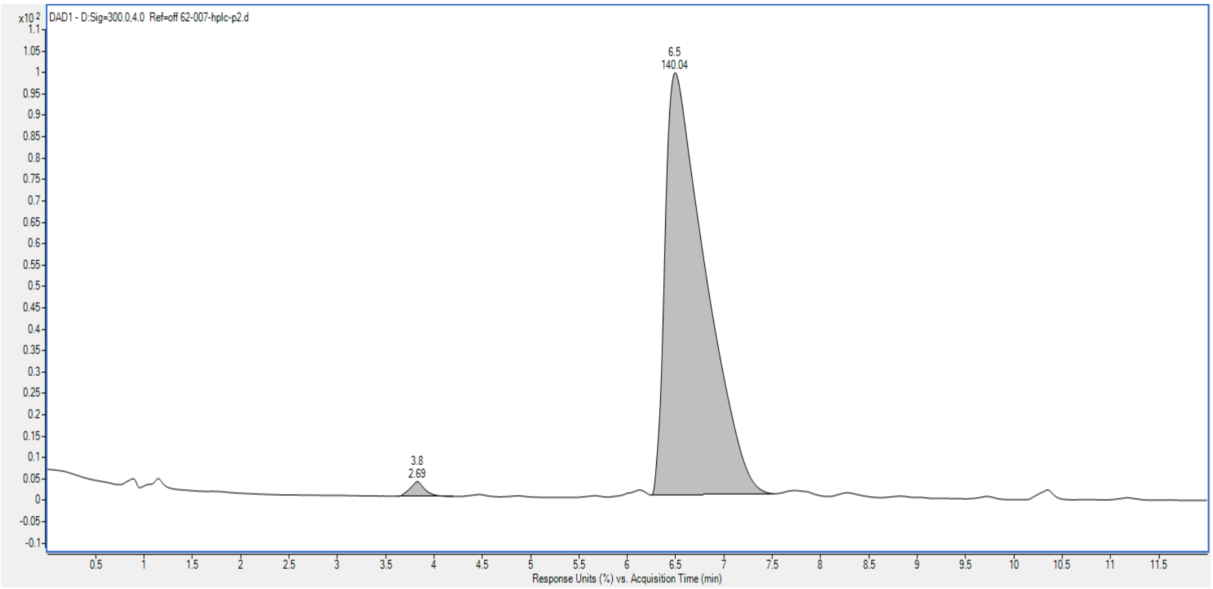


**Figure S23**. Analytical HPLC of **MKV3** ($t_R$ = 6.5 min)

**Table S1: MST analysis of compound binding to human ATP7A.**

| COMPOUND | BINDING AFFINITY (kD; nM) |
|---|---|
| MKV1 | 1831 ± 151 |
| MKV2 | Not detected |
| MKV3 | 219 ± 14 |

Microscale scale thermophoresis (MST) was performed using detergent-free lysates prepared from HEK293 cells expressing GFP-tagged human ATP7A.

**Table S2: Predicted MKV3-interacting residues within *Xt*ATP7B and correponding residues in human ATP7A and ATP7B**

| ***Xt*ATP7B** | **ATP7A** | **ATP7B** |
|---|---|---|
| H640 | H643 | H643 |
| E643 | E646 | E646 |
| W647 | W650 | W650 |
| Y710 | Y730 | Y713 |
| Q714 | Q734 | Q717 |
| N725 | N745 | N728 |
| R775 | R795 | R778 |
| E778 | E798 | E781 |
| H779 | H799 | H782 |
| K782 | K802 | K785 |
| P909 | P929 | P912 |
| I910 | I930 | I913 |
| Q911 | Q931 | Q914 |
| Q912 | Q932 | Q915 |
| D915 | D935 | D918 |
| **E1010** | **E1030** | **K1013** |
